# Breeding alters females’ social positions by changing dominance dynamics

**DOI:** 10.1101/2023.09.20.558583

**Authors:** Tobit Dehnen, Brendah Nyaguthii, Wismer Cherono, Neeltje J. Boogert, Damien R. Farine

**Author notes:** **Author for correspondence:** Tobit Dehnen The Farmhouse, Centre for Ecology and Conservation, University of Exeter, Penryn Campus, Treliever Road, Penryn, Cornwall, TR10 9FE, UK.

## Abstract

Agonistic and affiliative interactions with group members dictate individuals’ access to resources, and investment in competing for resources is often traded off with other needs. For example, reproductive investment can reduce body condition and, thereby, an individual’s ability to win future agonistic interactions. However, group members may also alter their behaviour towards reproductive individuals, such as becoming more or less aggressive. Here, we investigated the social consequences of reproduction in female vulturine guineafowl *Acryllium vulturinum*, a plural breeder in which females disperse and are subordinate to males. We found opposing patterns for breeders’ within- and between-sex dominance interactions from before to after breeding. Specifically, while breeding females became far less likely to win dominance interactions with non-breeding females after breeding than before breeding, they received considerably fewer male aggressions than non-breeding females after breeding. Using agent-based models, we then show that hierarchies inferred using the Percolation and Conductance method are robust to such variation in interaction rates, while other common methods may be prone to systematically overestimating or underestimating particular individuals’ hierarchy positions. Our study highlights reproduction as a driver of dominance dynamics, and illustrates how the study of dominance may benefit from considering variation in interaction rates.

## Introduction

In many group-living species individuals occupy particular social positions, defined by their associations as well as agonistic and affiliative relationships with group members [1,2]. Dominance relationships for example are a prominent feature of many animal societies [3], with individuals’ relative dominance dictating access to resources, such as reproductive opportunities [4]. However, investment in reproduction must also be traded off with investment elsewhere [5–7], such as maintaining body condition or engaging in social interactions. Given that engaging in, and escalating, agonistic interactions is costly [8,9], individuals investing heavily in reproduction may become less likely to win agonistic interactions with group members.

While reproduction may detrimentally influence outcomes of individuals’ intrasexual dominance interactions, reproducing may, conversely, positively alter individuals’ interactions with the opposite sex. Reproducing individuals likely care for the offspring of at least one opposite-sex group member (or more, if there is high within-group relatedness among the opposite sex or if the reproducing individual has mated multiply). Such opposite-sex group members should benefit from the survival of related juveniles [10], and thus also from the survival of their carer. Reproducing individuals should therefore become more valuable to some members of the opposite sex, which may thus alter their interactions—e.g. being more tolerant, or engaging in more affiliative or cooperative interactions—with such reproducing individuals. For example, in barbary macaques *Macaca sylvanus* breeding females receive more male grooming relative to non-breeding females, despite these females not increasing the grooming they give to other males [11]. Accordingly, reproducing may modulate both intrasexual and intersexual social interactions of both reproducing and non-reproducing individuals.

One situation where the influence of reproduction on intrasexual and intersexual interactions can be studied is sex-stratified hierarchies [12] when primarily the subordinate sex invests in reproduction. In such scenarios, reproduction-related changes in intrasexual agonistic interactions should be largely driven by individuals’ relative competitive abilities. Additionally, changes in intersexual agonistic interactions should be driven by the costs and benefits to dominant-sex individuals of aggressing particular subordinate, opposite-sex group members—which may change with such subordinate-sex group members’ reproductive status, as outlined above. In most mammal systems, the dominant sex either disperses (as in many primates, e.g. chacma baboons *Papio ursinus* [12] and crested macaques *Macaca nigra* [13]) or invests more heavily in reproduction (e.g. as in spotted hyenas [14]). In species where the dominant sex disperses, subordinates are philopatric and thus likely have within-group relatives and inherit their parents’ social relationships, maintaining these over their lifetimes ([15], e.g. spotted hyenas [16]; African elephants *Loxodonta africana* [17]; rhesus macaques *Macaca mulatta* [18,19]). Meanwhile, in species where the dominant sex invests in reproduction, studying reproduction-related changes in tolerance by opposite-sex individuals is largely redundant. Thus, it may be difficult to simultaneously investigate the effects of reproduction on intrasexual and intersexual agonistic interactions in many mammalian systems. In contrast, in birds females often i) are the dispersing sex [20]; ii) are subordinate to males (as in many mammals) given their smaller body size; and iii) invest heavily in reproduction [21].

Here we examine the temporal dynamics of the outcomes of intrasexual dominance interactions, as well as the rate of male-to-female aggressions, in relation to females’ reproductive status in vulturine guineafowl *Acryllium vulturinum*. This is a species in which females, the subordinate sex [22], both invest highly in reproduction [23] and disperse [24], while the dominant sex is philopatric. This means that (i) the body condition of individuals of the subordinate sex can be expected to change over time as a consequence of reproductive investment in egg production and incubation [23] (as in other Galliformes, e.g. female ring-necked pheasants *Phasianus colchicus* lose 19% of their body condition, primarily during incubation [25]); (ii) individuals of the subordinate sex can be expected to have no or limited within-group relationships prior to integrating into a new group. Accordingly, changes in females’ intrasexual dominance interactions surrounding breeding should primarily be driven by their relative competitive abilities. Meanwhile, as (iii) adult males are socially dominant to females at all times [22], changes in the rates of intersexual aggression received by females should predominantly be driven by males’ behaviour in response to females’ reproductive status.

We specifically investigated outcomes of dominance interactions among females, and the rates of male aggression towards individual females, both before and after breeding in relation to females’ reproductive status. This enabled us to quantify the short-term effects of reproduction in the subordinate, dispersing sex that occurred via both female-female interactions and male-female aggressions. We predicted that breeders (relative to non-breeding females) would be less likely to win female-female dominance interactions against non-breeders after, but not before, breeding, and that breeders (relative to non-breeding females) would face less male aggression than non-breeders after, but not before, breeding. Lastly, we used agent-based models to assess whether variation in interaction rates can cause particular individuals’ hierarchy positions to be systematically overestimated or underestimated when using common dominance hierarchy inference methods.

## Methods

### Data collection

We studied a habituated vulturine guineafowl social group at Mpala Research Centre, Kenya, in a population that has been studied since 2016. Breeding and interaction data used in this study were collected between 4^th^ March 2020 and 23^rd^ September 2022. The site comprises a mix of savannah and dense, dry woodland habitat, with glades (open, grassy areas) spread throughout the study site. Our data collection is limited to this group, and this period, because these birds live (and reproduce) predominately within a fenced compound, enabling us to follow them on foot, and because an extended drought starting in late 2020 meant that there was no subsequent breeding.

### Study species

Vulturine guineafowl live in stable social groups that range in size from 13 to 65 adult individuals [26] in dry woodland habitats in East Africa. Groups comprise more males than females and are characterized by steep dominance hierarchies with all adult males dominant to females [22]. Males’ social influence over females’ resource access is thus likely to be considerable. Groups tend to spend early mornings and late afternoons foraging and interacting on glades—typically feeding on grass leaves, roots, bulbs and seeds, as well as berries and small invertebrates [27,28]. Such prolonged exposure in open areas makes it easy to observe aggressive and submissive behaviours among group members [22,29].

Seasonality at our study site is characterized by wet and dry periods, with breeding taking place during the wet periods [28]. Breeding starts with male-female pairs forming and moving separately from the remaining social group, with the male typically mate-guarding the female against other males [23,26]. While many females may form temporary pairs, only a portion of adult female group members typically attempt to breed in any given season [23]. Females that breed lay eggs in a well-hidden scrape on the ground on consecutive days, until they reach a clutch size of up to 15 eggs, although clutch sizes of eight to ten eggs are more typical in our population [23]. The female then incubates the nest for approximately 23 days [27]. We have no observations of females receiving help during incubation (females of the closely related helmeted guineafowl *Numida meleagris* incubate alone [30]), nor do females take any extensive foraging breaks during the incubation period. Meanwhile, males paired with incubating females re-join the social group and often re-pair with another female [23]. High rates of nest predation in this ground-nesting species mean that only some females that attempt to breed reach incubation and hatching [23]. After breeding, the social group re-forms as females return with their broods within days of hatching [26] or their nest being depredated. Distinct subsets of male group members then help females raise their offspring, largely doing so by calling chicks to food items and covering chicks under their wings [23]. Once the group has reformed, dominance interactions among group members can readily be observed.

### Data collection: breeding

During breeding seasons, we monitored the breeding attempts of adult females in the social group. We endeavoured to follow paired females to determine the location of nests. We then monitored located nests to determine the number of eggs therein and the breeding stage reached. For the purposes of this study, we defined individuals that successfully hatched chicks as ‘breeders’, given that these individuals produced a clutch of eggs and incubated these for approximately 23 days [27]. We defined ‘non-breeders’ as those individuals that did not attempt to breed or those whose nests failed prior to hatching, given that this could happen after only few eggs were laid and without incubation. Accordingly, in our dataset, two females whose nests were predated—the first during the laying stage (season one) and the other after approximately one week of incubation (season two)—were categorized as non-breeders. While we acknowledge that this is a somewhat arbitrary categorisation along a continuum of investment in breeding, our categorisation does present a clear and obvious distinction. Additionally, while females whose nests failed did invest in breeding, such females both invested to a lesser degree and returned to the group after only a short absence—while females who bred successfully were absent for the entirety of incubation, over three weeks.

For each breeding female in each season, we identified the first and last date of observed breeding behaviour (laying, incubating or hatching). However, some females were absent for periods equivalent in duration to incubation and subsequently returned to the group with chicks, yet their nests were never found. We treated these individuals as breeders. For each female we therefore subtracted one month (23 days incubation + eight eggs laid on consecutive days) from the hatch date to estimate the start of their breeding behaviour. If this estimate was prior to the female’s first known breeding behaviour, the estimate was used as the start of breeding instead. We then defined the “breeding period” for the social group in that season as the period of time between the first and last date of observed breeding behaviour from individuals defined as breeders in that group and season. Note that we do not consider post-hatching parental care behaviour as breeding behaviour. This is because the precocial vulturine guineafowl chicks are relatively independent from a young age, and females return to the social group almost immediately upon chicks hatching—after which a considerable portion of offspring care is provided by male group members [23]. Therefore, our breeding-season analysis considers dominance interactions before breeding females leave the social group and once they return.

To get an approximation of female investment in egg production, we weighed eggs from three females’ clutches during season two using Kenex Eternity 100 scales. We then combined this egg weight data with female body weight measurements from trapping data. Specifically, we estimated female investment in egg production as a percentage of body mass via: 100 × no. eggs (eight or ten) × median egg weight / median female body weight. While this is unlikely to reflect all breeders’ investments due to differences in clutch size, and egg and body weights, it nonetheless provides an indication of female investment in egg production.

### Data collection: dominance interactions

We used all-occurrence sampling [31] to record dyadic dominance interactions (both aggressive and submissive) among group members, typically during mornings and late afternoons when birds are most active, and also recorded the group composition and observation duration for each session. As vulturine guineafowl readily form subgroups [29] a new observation session was started when the group composition changed. Using this approach, we have established a large interactions dataset comprising different types of dominance interactions across years and social groups [22,29]. We considered the actors of aggressive interactions and the recipients of submissive interactions to be the ‘winners’, and the recipients of aggressive interactions and the actors of submissive interactions to be the ‘losers’. We used subsets of this long-term interaction dataset in all analyses.

### Construction of the datasets: female-female interactions surrounding the breeding season

We used the long-term interactions dataset for each season in the breeding-season analysis, as follows. First, we created a 22-week subset that was 11 weeks before and 11 weeks after the halfway point of the ‘breeding period’ (defined above). We then removed any interactions from this subset that occurred during the breeding period and/or involved males, splitting the season’s data into two approximately eight-week subsets of female-female dominance interaction data: one before and one after breeding took place (see Figure S1). We also removed any interactions involving females that hatched after that season’s start date (i.e. hatched during that season). We then merged all data subsets into one dataset containing the data of all pre- and post-breeding periods of both seasons. We thus obtained a winner-loser dataset containing interactions among and within breeding and non-breeding females from both before breeding and after breeding for both seasons (see Figure S1 for timeline).

### Construction of the datasets: male-female aggressions surrounding the breeding season

We used interaction data from the same time periods as the breeding-season analysis, above. For each data collection session, we calculated the number of male aggressions experienced by each female, as well as the number of males present and the session duration. The latter two measures were multiplied to produce a male-hours variable capturing the exposure of females to males during each data collection session (and thus opportunity for females present to be aggressed by males). We then summed both the number of male aggressions and the male-hours from all the data collection sessions for each female in each period in each season. This produced a total number of male aggressions received and total male-hours for each female in each period, i.e. before and after breeding, in each season.

### Analytical approaches

Studying dominance dynamics using longitudinal or repeated static hierarchies can be difficult. The interdependence of individuals’ inferred hierarchy positions [32] mean that individuals’ hierarchy positions, and associated errors therein, are highly correlated. For example, if an individual is ranked incorrectly then, by definition, other group members will also be ranked incorrectly. Additionally, measurement uncertainty is often not explicitly considered and the time steps at which hierarchies are updated can influence hierarchy dynamics [33]. Furthermore, passive demographic processes can drive hierarchy dynamics [34], which we are not interested in here. Accordingly, many of our analyses relied on interaction-level data and generalized linear mixed effects models, as used in previous studies of contest outcomes [35,36].

Throughout, we assessed model fits using the R package DHARMa [37], and assessed statistical significance of model fixed effects using type-II (for models with no interaction terms) and type-III (for models with interaction terms) Wald χ2-tests in the R package car [38] unless specified otherwise. We conducted all statistical analyses using R version 4.1.2 [39] (aside from the ASReml models, which were run in R version 4.3.0) and considered P < 0.05 as significant. We used the R packages sjPlot [40], ggplot2 [41] igraph [42] and cowplot [43] to create figures.

### Dyadic-interaction model overview

In our interaction-level datasets there was one winner and one loser for every interaction. As winner-loser data consist of two interacting individuals but only one outcome, we randomly assigned one individual as the ‘focal’ (focal_ID) and one as the ‘interactor’ (interactor_ID) for each interaction. In such dyadic contest data, individual a winning against individual b could be represented as either focal_a_ beats (1) interactor_b_, or focal_b_ loses (0) to interactor_a_. The outcome (1/0) is therefore entirely dependent on the allocation of the two individuals to ‘focal’ and ‘interactor’ roles, which is important in datasets comprising multiple observations per individual (see [44]), as is the case for our interaction data. In our analyses using interaction-level data, we therefore associated each observation (interaction) with two levels (focal_ID and interactor_ID) of a single, additive random effect to model focal and interactor identities.

### Breeding-season analysis

To assess whether females’ probabilities of winning intrasexual dominance interactions changed as a consequence of breeding, we fitted a generalized linear mixed-effects model to the dataset of female-female interactions surrounding the breeding season using the glmer function of the R package lmerMultiMember [45]. This package provides a wrapper for the glmer/lmer functions of the R package lme4 [46], and allows the modelling of focal and interactor identities as a single, additive random effect for winner-loser data with multiple observations per individual. This replicates the random effects structure of ASReml models in similar analyses [36,44]. We chose to use lmerMultiMember over ASReml software as lmerMultiMember allows for binomial error structures (which are correct for our data, while ASReml allows only Gaussian). As the lmerMultiMember package is relatively new, we replicated the breeding-season analysis (see below) using the R package ASReml, which has been widely used for modelling dyadic interactions for over a decade [44]. Our ASReml replicate confirms that model outputs are qualitatively the same regardless of the R package used (see supplementary materials).

We fitted focal_won, whether the focal won (1) or lost (0) the dominance interaction, as the response variable and used a binomial error structure. Fixed effects included: period, whether the interaction took place before or after breeding; dyadic_breeding_contrast, a three-way categorisation of the focal and interactor combined breeding status (either focal = breeder & interactor = non-breeder, focal = non-breeder & interactor = breeder, or focal & interactor are of the same breeding status). Under the hypothesis that breeders are less likely to win against non-breeders—and vice versa for non-breeders against breeders—after breeding, the interaction term between period and dyadic_breeding_contrast is predicted to be significant. Specifically, predicted values are predicted to diverge for breeding and non-breeding individuals after breeding, such that non-breeding individuals are more likely to win against breeding individuals. We also included season, i.e. which season the interaction took place in (season one or two), as a fixed effect (due to having only two levels). Random effects included: focal_ID and interactor_ID (defined above) as a single, additive random effect as outlined above; and Dyad_ID, an ID unique to the interacting dyad (irrespective of focal and interactor allocation) as a further random effect. We tested the significance of random effects using likelihood ratio tests comparing models with and without the random effect in question.

### Male-female analysis

To assess whether the level of male aggression experienced by females varied according to breeding status and period, we fitted a generalized linear mixed-effects model to the dataset on male-female aggressions surrounding the breeding season. We fitted the number of male aggressions received (num_aggro) as the response variable. To account for opportunity for females to be aggressed by males, we log-transformed the num_male_hours variable (which captures females’ exposure to males) to produce the variable log_num_males_hours and fitted this as an offset. We fitted both period and the female’s breeding status (breeder_status), as well as the interaction term between them, as fixed effects. As in other models, we wanted to control for season, which we fitted as a fixed effect due to having only two levels. We fitted the female’s identity (ID.original) as a random effect, given that each female contributed at least two datapoints. We fitted the initial model with a Poisson error distribution using the R package lme4 [46], but this suffered from overdispersion. We thus fitted a negative binomial model instead using the R package glmmTMB [47]. This model converged without problems and was used for statistical analysis.

### Influence of variation in interaction rates on hierarchy inference

Conventional hierarchy inference methods typically rely on an even distribution of individuals in interaction data. Subtle variation in dyadic interaction rates among group members could therefore influence hierarchy inference. We tested this using agent-based models as follows:

1) Generate a group comprising 16 males and 10 females, emulating sex ratios and group sizes seen in vulturine guineafowl [22,48].
2) Assign individuals a dominance value (representing all determinants of dominance in one value) from two normal distributions, such that males are dominant over all females, and assign females’ ‘real hierarchy positions’ as the order of their dominance values (for subsequent comparisons with their estimated hierarchy positions).
3) Randomly assign 50% of females as ‘breeding’ and 50% as ‘non-breeding’.
4) Simulate outcomes of dyadic dominance interactions via a probabilistic approach based on interacting individuals’ relative dominance values and a hierarchy steepness parameter. Thus, interaction outcomes directly reflect the dominance values (with additional noise).
5) Determine the outcome of individual interactions in two scenarios: (i) males tolerate, and thus engage in interactions with, breeding and non-breeding females equally, or (ii) males tolerate breeding females and redirect intersexual interactions towards non-breeding females. Additionally, given that females may evade potential interactions by associating with dominant group members, in this scenario breeding females engage in only half as many intrasexual interactions with non-breeding females (for illustrated aggression network examples see Figure S7).
6) For each scenario, infer the dominance hierarchy (via one of the methods outlined below) from the simulated interaction outcomes, extract among-female rank orders from the inferred hierarchy, then subtract each females’ real hierarchy position (see step two) from its inferred hierarchy position, and calculate the mean difference for both breeding and non-breeding females. For example, if five breeding females’ inferred hierarchy positions minus real hierarchy positions were 1, 2, 0, 1, 1, then this would result in a mean of (1 + 2 + 0 + 1 + 1) / 5, i.e. one, and suggest that breeding females’ hierarchy positions were estimated to be higher than their real hierarchy positions.

We repeated the above processes 1000 times and then plotted density distributions, as well as 15, 50 and 85% quantiles, of outputs. This allowed us to compare how well females’ inferred hierarchy positions reflected their real hierarchy positions, and how this differed between the two scenarios. Under the hypothesis of no systematic overestimation or underestimation of individuals’ hierarchy positions, output values should centre equally around zero. We repeated this procedure for three common dominance hierarchy inference methods: randomised Elo ratings [32], the I&SI method [49] and the Percolation and Conductance approach [50] (see Supplementary Materials for full details).

## Results

### Egg investment

Vulturine guineafowl eggs (n_eggs_ = 19; n_females_ = 3) weighed a median of 46g (range: 43–52g), with the median body weight of a female (n_weights_ = 882; n_females_ = 533) being 1380g (range: 1030–1840g). Given these estimates of egg and female body weights, and clutch sizes of eight to ten eggs [23], egg production represented approximately 27–33% of females’ body weight and thus a substantial investment to females alongside incubation effort.

### Breeding-season analysis

Our breeding-season analysis involved 1325 female-female interactions from a total of 116 unique dyads (see Table 1 for season- and period-level details). We found no evidence for either period (χ^2^_1_ = 0.142, P = 0.706) or dyadic breeding contrast (χ^2^_2_ = 1.848, P = 0.397) to affect focal winning probability in our breeding-season analysis. Likewise, we found no significant difference between focal individuals’ winning probabilities between seasons (χ^2^_1_ = 0.0525, P = 0.819). However, as predicted, there was a significant interaction between period and dyadic breeding contrast (χ^2^_2_ = 33.309, P < 0.001). Specifically, non-breeders experienced an increased probability of winning against breeders after the breeding period, while breeders experienced a corresponding decreased probability of winning against non-breeders after the breeding period. Meanwhile, breeders and non-breeders appeared to be equally likely to win before breeding (Figure 1).

**Table 1.**
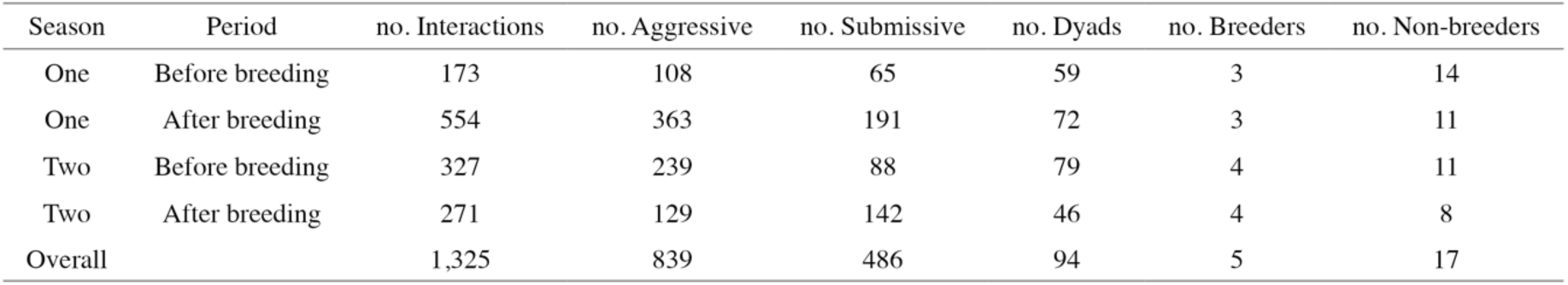
Summary of female-female dominance data from each season and period therein, used in the breeding-season analysis.

**Figure 1.**
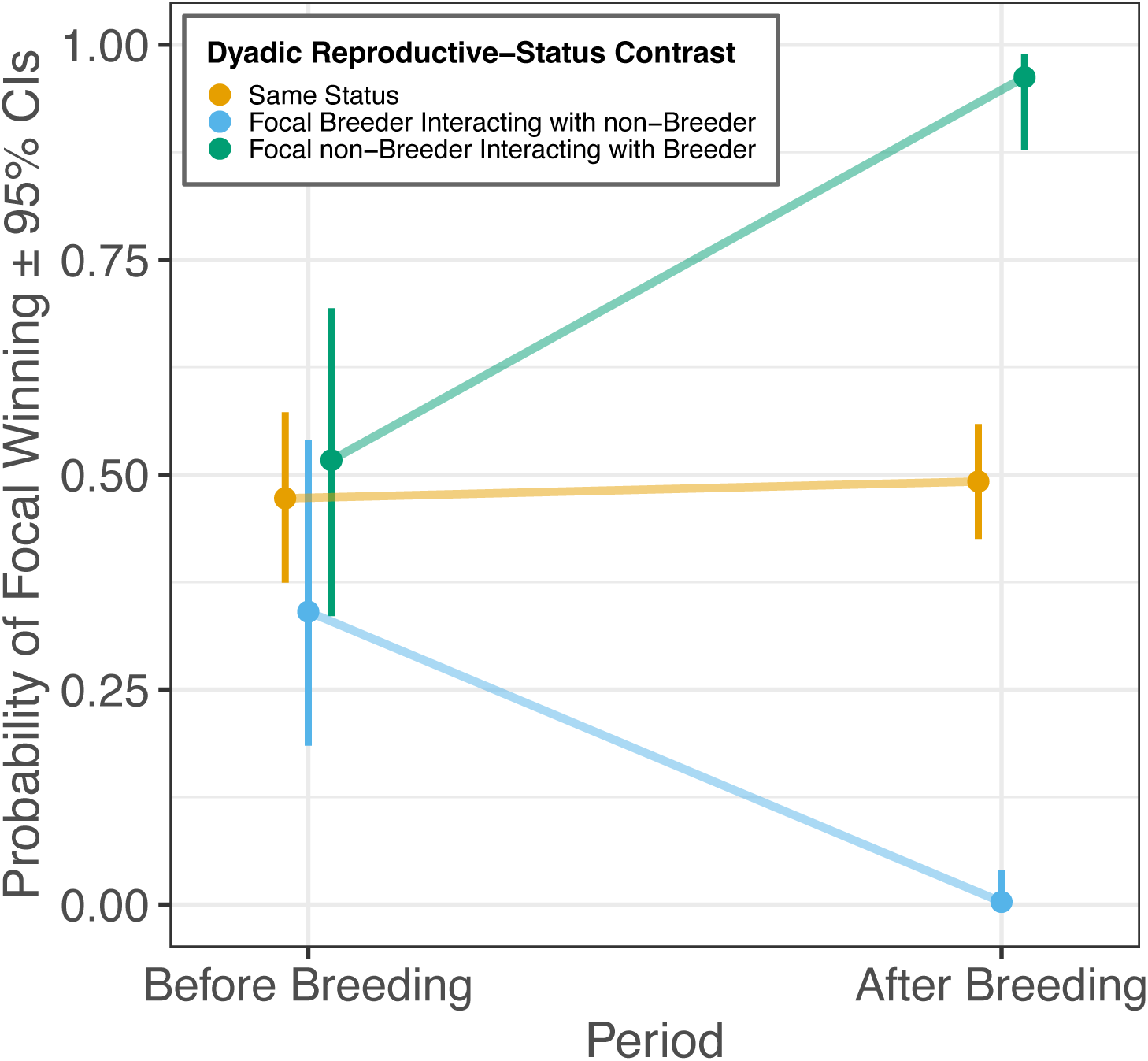
Breeding females are more likely to lose interactions with non-breeding females after, but not before, breeding. Dyads where the focal and interacting individual are of the same breeding status (yellow points) comprise both breeder–breeder and non-breeder–non-breeder interactions.

Only females observed being aggressive before a season started were included.

Testing of random effects in the breeding-season analysis revealed between-individual differences in competitive ability. Specifically, the model including the additive term of focal ID and interactor ID outperformed the model excluding this term (χ^2^ = 751.524, P < 0.001), indicating that individuals differed in competitive ability beyond their breeding status. Adding dyad ID as a random effect did not further improve the model beyond the additive effect of the interacting individuals’ identities (χ^2^ = 0.082, P = 0.774) (Table 2). This result suggests that interactions among females in the breeding-season analysis dataset are highly transitive.

**Table 2.** Likelihood ratio tests comparing breeding-season analysis models that differ only in random effects structure.

| Model random effect structure | *AIC* | *Log-Likelihood* | $\chi^2$ | *p value* |
| --- | --- | --- | --- | --- |
| season | 1,779.34 | -882.67 |  |  |
| season + additive individual IDs | 1,029.82 | -506.91 | 751.52 | **<0.001** |
| season + additive individual IDs + dyad ID | 1,031.74 | -506.87 | 0.08 | 0.774 |

Each model is compared to the model above, with models in which the additional random effect significantly improved model performance highlighted in bold.

### Male-female analysis

We found that the number of male aggressions females faced differed between seasons (χ^2^_1_ = 56.780, P < 0.001), being higher in season two. We found no evidence for a difference in aggression faced between breeding and non-breeding females across all periods and seasons (χ^2^_1_ = 0.270, P = 0.604). However, while the number of male aggressions faced by females was the same for breeders and non-breeders before breeding, the level of aggression faced diverged after breeding according to females’ breeding status. Specifically, non-breeding females faced considerably more male aggression than breeding females after breeding (Figure 2), as highlighted by the significant interaction between period and breeder status (χ^2^_1_ = 10.709, P = 0.001). This enhanced aggression faced by non-breeding females after the breeding period likely largely drove the greater aggression faced by females overall after breeding relative to before breeding (χ^2^_1_ = 80.110, P < 0.001). However, the difference in aggression received before versus after breeding could also be a product of sampling method (see discussion).

**Figure 2.**
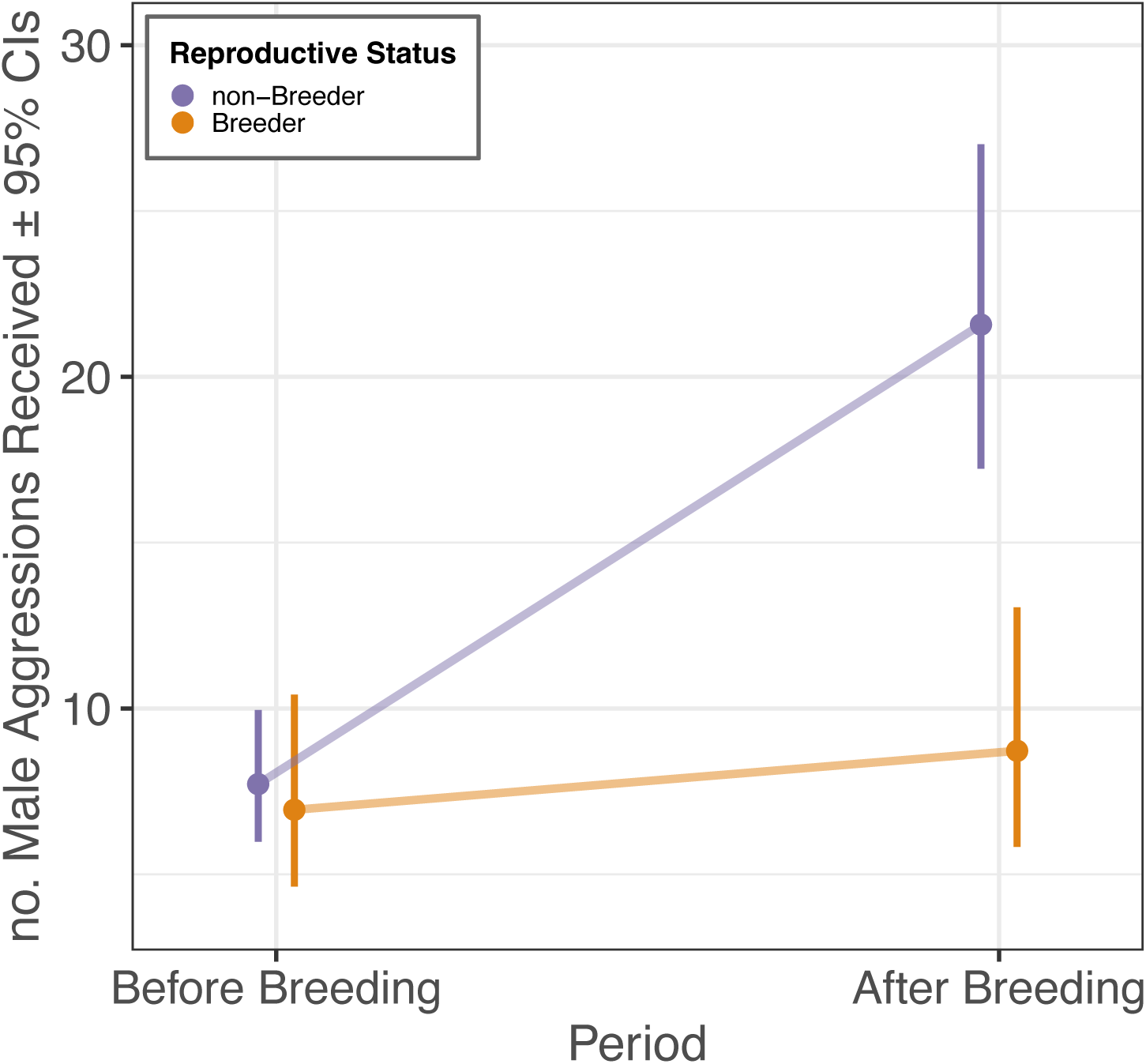
Breeding females receive less aggression from males than non-breeders after, but not before, breeding. The number of aggressions received by females was corrected to account for females’ exposure to males, calculated as log(total number of male-hours a female was exposed to), using an offset term in the model.

### Influence of variation in interaction rates on hierarchy inference

In our agent-based model where males had no biased tolerance towards certain females, we found that individuals’ inferred hierarchy positions were consistent with their real hierarchy positions in both breeding and non-breeding females across all hierarchy inference methods tested (see blue distributions in Figure 3). However, when males were tolerant of breeding females only, breeding females’ hierarchy positions were systematically estimated as higher than their real hierarchy positions (while those of non-breeding females were correspondingly estimated as lower) when inferred using randomised Elo ratings or the I&SI method (see yellow distributions in Figure 3A–D). Exploration of the parameter space suggested that, while the degree of overestimation and underestimation varied with sex ratio, hierarchy steepness, data depth (i.e. the number of interactions ‘observed’ while dyadic ratios are maintained) and the proportion of females that were allocated as ‘breeders’, the effect was always present in the biased scenario when hierarchies were inferred using randomised Elo rating or I&SI approaches. Furthermore, when comparing females’ inferred vs real hierarchy positions across the entire group—rather than only within females—the degree of overestimation of breeding females’ hierarchy positions in the scenario of biased male tolerance was more pronounced (see Figure S8). This occurred because breeding females were sometimes inferred as having higher hierarchy positions than some males. In contrast, when inferred using the Percolation and Conductance method [50], the inferred hierarchy positions of breeding and non-breeding females—on average—matched their real hierarchy positions in both scenarios (Figure 3E,F), with minimal systematic overestimation or underestimation across all parameters tested (see Supplementary Materials).

**Figure 3.**
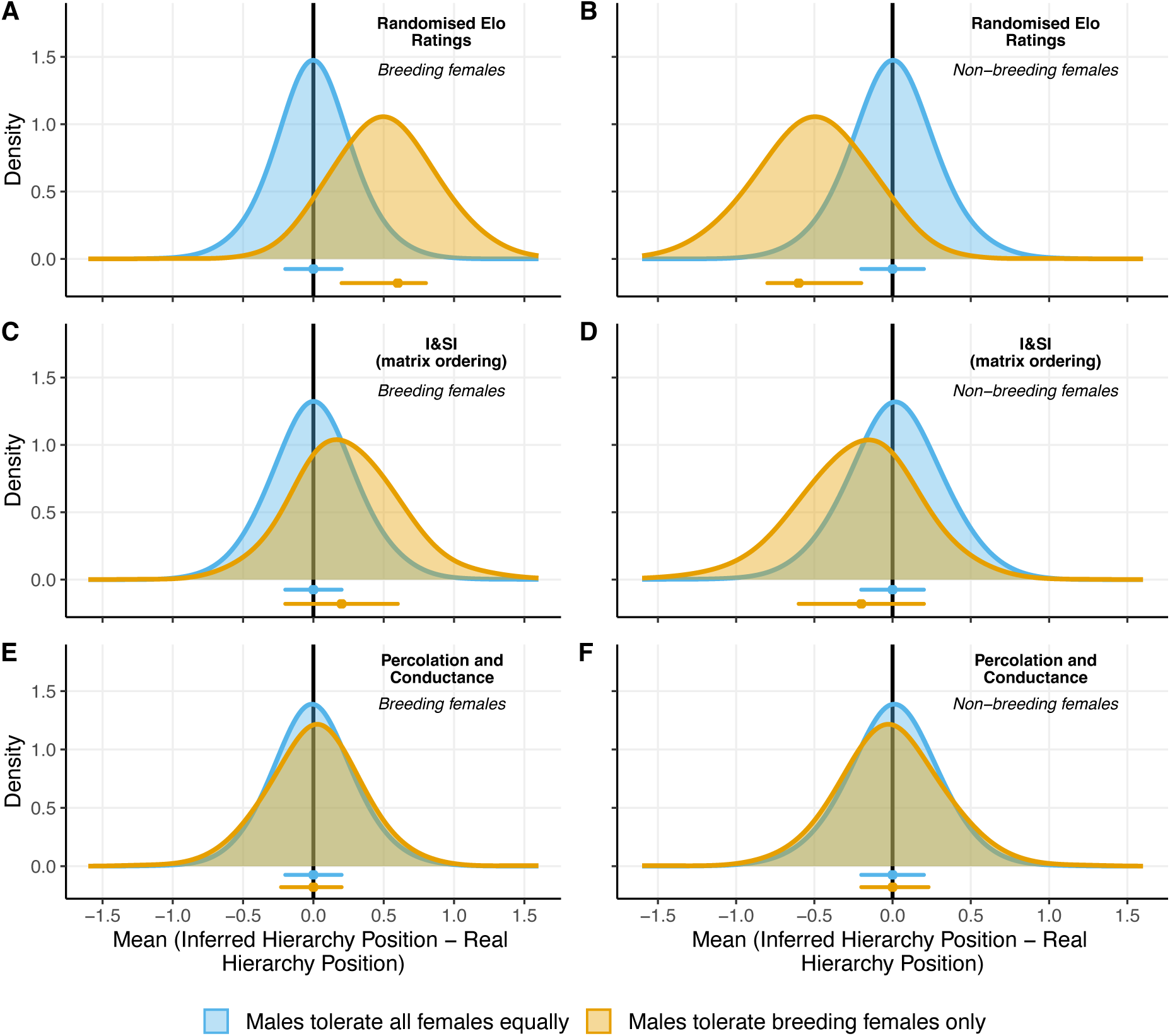
Agent-based model results showing the consequences of changes in both patterns of dominance interactions as well as inference method on hierarchy inference. When using common approaches for dominance hierarchy inference, females that were involved in fewer intersexual and intrasexual interactions were estimated to be positioned higher in the inferred dominance hierarchy relative to their real hierarchy positions (A,C). In contrast, females who received more intersexual aggression were estimated to have disproportionately lower hierarchy positions (B,D). Meanwhile, we observe no such difference in inferred versus real hierarchy positions when using the Percolation and Conductance method (E,F). Results are based on 1000 simulations of interactions among 16 males and 10 females from two scenarios: either males tolerate all females equally (blue), or males preferentially tolerate ‘breeding’ females (yellow). The latter scenario is characterised by males redirecting intersexual interactions from breeding towards non-breeding females, and breeding females engaging in half as many interactions with non-breeding females as in the unbiased scenario. Points and horizontal lines represent the medians and 15-85% quantile ranges, respectively. Black vertical lines represent the expected value, zero, under the assumption that inferred hierarchy positions reflect real hierarchy positions. Real hierarchy positions here are the order of individuals’ hardcoded dominance values, used to determine the outcomes of all interactions.

## Discussion

In many group-living species reproductive success is skewed towards dominant individuals [4,51], with dominant individuals breeding more often, producing larger offspring or producing more offspring per reproductive event [52]. However, reproduction can also be a predictor of future dominance outcomes. Our analysis of female-female dominance interactions surrounding breeding seasons revealed that, while breeders had an equal probability of winning dominance interactions with non-breeders before breeding, this reduced considerably after breeding (Figure 1) and remained so for at least three months (see supplementary materials). We also found no evidence that breeders’ winning probability declined further between two post-breeding periods (Figure S2; Table S1). This suggests that the observed pattern of breeding and associated winning probability dynamics were not driven by individuals in decline choosing to breed (akin to terminal investment [53]), but that the reduced winning probability of breeders was a consequence of reproducing. Furthermore, examination of random effects in our breeding-season analysis suggests that dominance in female vulturine guineafowl was highly transitive, and that females differed in competitive ability beyond their breeding status alone (Table 2). Our analysis of male-female interactions surrounding breeding seasons then showed that breeding females were the targets of less intersexual, male aggressive interactions when compared to non-breeding females after, but not before, breeding. While limited to five breeding and 17 non-breeding females across two seasons, our data suggest that breeding can have opposing consequences for females’ relationships with female vs. male group members.

### Female reproduction and short-term intrasexual interaction outcomes

Breeding female vulturine guineafowl invest considerably in reproduction by producing a clutch of eggs (constituting ∼30% of their body mass) after a harsh dry season [23], during which groups’ home ranges expand in the pursuit of resources for survival [28]. Additionally, females incubate their clutch solitarily and seldom leave the nest. Meanwhile, non-breeding females may continue to forage during the resource-rich wet season [28]. We therefore stipulated that reduced condition in breeding individuals drives the reduction in breeding females’ winning probabilities. Our analysis of interactions surrounding trapping events outside of the breeding season however suggests that, while structural size has some influence on interacting outcomes, body condition appears to play a lesser role (Figure S6). Nevertheless, at extreme differences in body condition the individual in vastly superior condition always won (see Figure S6). If reproductive investment results in such extreme differences in body condition among breeders and non-breeders, condition could indeed be the proximate mechanism driving the observed divergence in breeders’ and non-breeders’ winning probabilities after breeding. A non-mutually exclusive, alternative possible driver of breeders’ post-breeding decline in winning probability is that breeding females are effectively removed from the social group for at least the duration of incubation, around one month (see horizontal lines in Figure S1). In many species, recent dyadic interaction outcomes determine individuals’ behaviour in future interactions within that dyad [54]. This effect relies on memory and so can fade over timespans similar to the incubation period of vulturine guineafowl [54]. Breeders’ absences from the social group may thus ‘release the shackles’ on prior subordinates—even in the absence of changes to females’ intrinsic attributes such as body condition, simply by breeders’ temporary removal. In spotted hyenas for example, females returning to a social group after being absent for over six months received severe aggression and suffered reduced dominance relative to their hierarchy positions prior to leaving [55]. Similarly, experimental removal and reintroductions of individual monk parakeets *Myiopsitta monachus* highlight that temporary, eight day absences can cause returning individuals to receive enhanced aggression and consequently drop down the hierarchy [56]. Accordingly, elucidating the mechanism driving the relationship between breeding status and interaction outcomes in female vulturine guineafowl will require experimental manipulation, e.g. through supplementary feeding or experimental removals, or the fine-scale tracking of individuals’ condition.

How and why reproduction influences interaction outcomes likely depends on how reproduction alters breeders’ and non-breeders’ determinants of dominance [54], which are dictated by species’ reproductive biology and the context of competitive interactions. For example, breeders should be more likely to win agonistic interactions in species where a) individuals frequently compete for monopolisable resources that represent tangible fitness benefits to breeders and b) physical reproductive investment is relatively low. In female parasitoid wasps *Goniozus nephantidis* and *Eupelmus vuilleti*, contests for access to hosts within which to lay their eggs are won by the individual that is carrying more eggs [57,58]. Similarly, reproductively experienced carrion roller beetles *Canthon cyanellus cyanellus* are more likely to win contests for food balls than naïve individuals [59]. As the respective authors suggest, these effects are likely driven by individuals’ asymmetric resource valuation, which arises from the difference in reproductive benefits of winning among. Conversely, breeders should be less likely to win agonistic interactions post-breeding in species where i) individuals rarely compete for monopolisable resources (or in which a single monopolisable resource represents a smaller fitness payoff), ii) breeders suffer reduced physical condition (i.e. intrinsic attributes [54]) as a consequence of short and intense reproductive investment (as is likely the case in many birds), and/or iii) individuals are temporarily absent from typically stable social groups [55,56] while reproducing, as a consequence of reduced recent interaction-outcome history with prior subordinates [54] or simply by many group members initiating agonistic interactions with returning individuals at an elevated rate. Reproductive status thus likely has diverse effects on interaction outcomes, and ultimately dominance relationships, among breeding and non-breeding individuals across species, with the pattern dictated by the species’ breeding biology.

### Female reproduction and short-term intersexual male-female aggressions

Our finding of lower levels of intersexual, male aggression experienced by breeding females after but not before breeding could be driven by several processes. Vulturine guineafowl societies are characterized by females dispersal and male philopatry [24], which likely lead to relatively high levels of relatedness among males of the same social group. Males are thus likely related to the offspring of one or more breeding females, which is reflected in their caring for juveniles that are unlikely to be their own offspring [23]. Being less aggressive, i.e. tolerant, towards mothers of such related offspring could thus carry indirect fitness benefits for males. The observed pattern could alternatively be driven by males directing more aggression towards non-breeding females. Due to female-biased dispersal [24], females that have dispersed and do not breed are unlikely to be closely related to any within-group chicks and thus incentivized to harm them [60]. Indeed, juvenile-directed female aggression is common among other species characterized by cooperative breeding [61] or dominance hierarchies [62], likely driven by female-female competition. Vulturine guineafowl males help a particular female raise her offspring, which, among other helping behaviours, includes protecting chicks from other adult group members [23]. The observed difference in aggression received by breeding and non-breeding females after breeding may therefore be driven by males protecting chicks by expressing enhanced aggression towards such non-breeding females. This latter explanation is in line with the model predicted values: after breeding the level of male aggression received increased for non-breeding females but remained relatively stable for breeding females (Figure 2). Accordingly, determining the mechanism driving breeding-status related differences in levels of male aggression received by breeding and non-breeding females after breeding is not possible with the present dataset. We advise caution when interpreting variation in predicted values between periods, i.e. before vs after breeding, as variation therein may be an artefact of the sampling method. Specifically, data were collected via all-occurrence sampling, for which observed dyadic rates likely decrease with increasing group size. As sampled subgroups are typically larger pre-breeding (when the entire social group is cohesive) than post-breeding (when broods and associated helpers move independently [23]), it is unclear whether the difference in pre-vs post-breeding predicted values reflect true rates or sampling artefacts. Nevertheless, within-period differences—between breeders and non-breeders either before or after breeding— should not be affected by any such sampling effects.

### Long-term consequences of reproduction

Kinship dynamics occur when individuals’ relatedness to their social group changes over time as a consequence of demographic processes and often operate in group-living species with pluralistic breeding [60,63]. For example, in chimpanzees *Pan troglodytes*, dispersal is strongly female-biased [64] meaning that, while initially low immediately after immigrating, females’ relatedness to group members increases over time via philopatry of male offspring [60]. As dispersal in vulturine guineafowl is likewise female-biased [24], as in most birds [20], similar kinship dynamics likely operate in vulturine guineafowl societies. Vulturine guineafowl groups also feature pluralistic breeding [23], which may be key to such kinship dynamics [60]. Relatedness of female vulturine guineafowl to their group members can thus be expected to increase over time as they reproduce. Given that individual kinship dynamics drive inclusive fitness payoffs of individuals’ helping and harming behaviour [63], female vulturine guineafowl may, as a consequence of generating philopatric sons, experience greater helping (e.g. tolerance, helper offspring care) and reduced harming (e.g. aggression) from their group members over time. Indeed, preliminary data (limited to the adult sons of a single female) suggest that male vulturine guineafowl exhibit role-reversed nepotism, whereby adult males are less aggressive towards their own mother than towards other female group members (Figure S5). Similar effects are found in spotted hyenas *Crocuta crocuta*, where cubs are tolerant of their (subordinate) fathers, towards which they direct less aggression than towards unrelated males [65]. Aside from kinship dynamics and associated benefits, females that breed successfully could also become more attractive to males, given that they have a track record of proven fecundity [66,67]. Our field observations suggest that forced copulations among vulturine guineafowl are uncommon (T. Dehnen, *personal observation*), meaning that females likely have considerable reproductive control. Males may thus exhibit enhanced tolerance towards previously breeding females that are potential mates. Reproduction may therefore have long-term consequences for vulturine guineafowl females’ social experience via multiple possible pathways. While examining long-term patterns of social integration and interactions in wild animal groups is not without its difficulties, we encourage future studies to investigate long-term transitions in individuals’ social ‘positions’ in relation to past breeding success.

By altering females’ relationships with males, breeding successfully could also have long-term consequences for females’ investment in intrasexual dominance. Attaining and maintaining social dominance is costly, as individuals have to invest time [68] and energy [8,9] in engaging in and escalating dominance interactions. However, females that have successfully bred may gain access to monopolisable resources via an alternate route—tolerance from related and unrelated males (as discussed above)—at least when in their presence. This may then allow previously breeding females to reduce investment in intrasexual dominance, and simultaneously allocate more resources to future reproduction. Thus, in the long-term, breeding may allow females to break free of a potential condition- and dominance-mediated negative feedback loop between resource access and reproduction. Meanwhile, females that lack such male tolerance should maximize investment in intrasexual dominance so that they may acquire resources to breed themselves and, ultimately, generate their own set of tolerant males. Females’ social relationships with male and female group members may thus change over time after entering the social group as a consequence of kinship dynamics (see Figure 4). Future work should investigate to what degree females both benefit from male tolerance at monopolisable food sources (e.g. quantifying food consumption) and invest in intrasexual dominance (e.g. receiving and initiating agonistic interactions), as a function of prior breeding success and the presence of particular male group members. Such effects could be studied in other group-living species with both male-biased dominance and agonistic interactions among females, as is likely the case in many mammals and birds.

**Figure 4.**
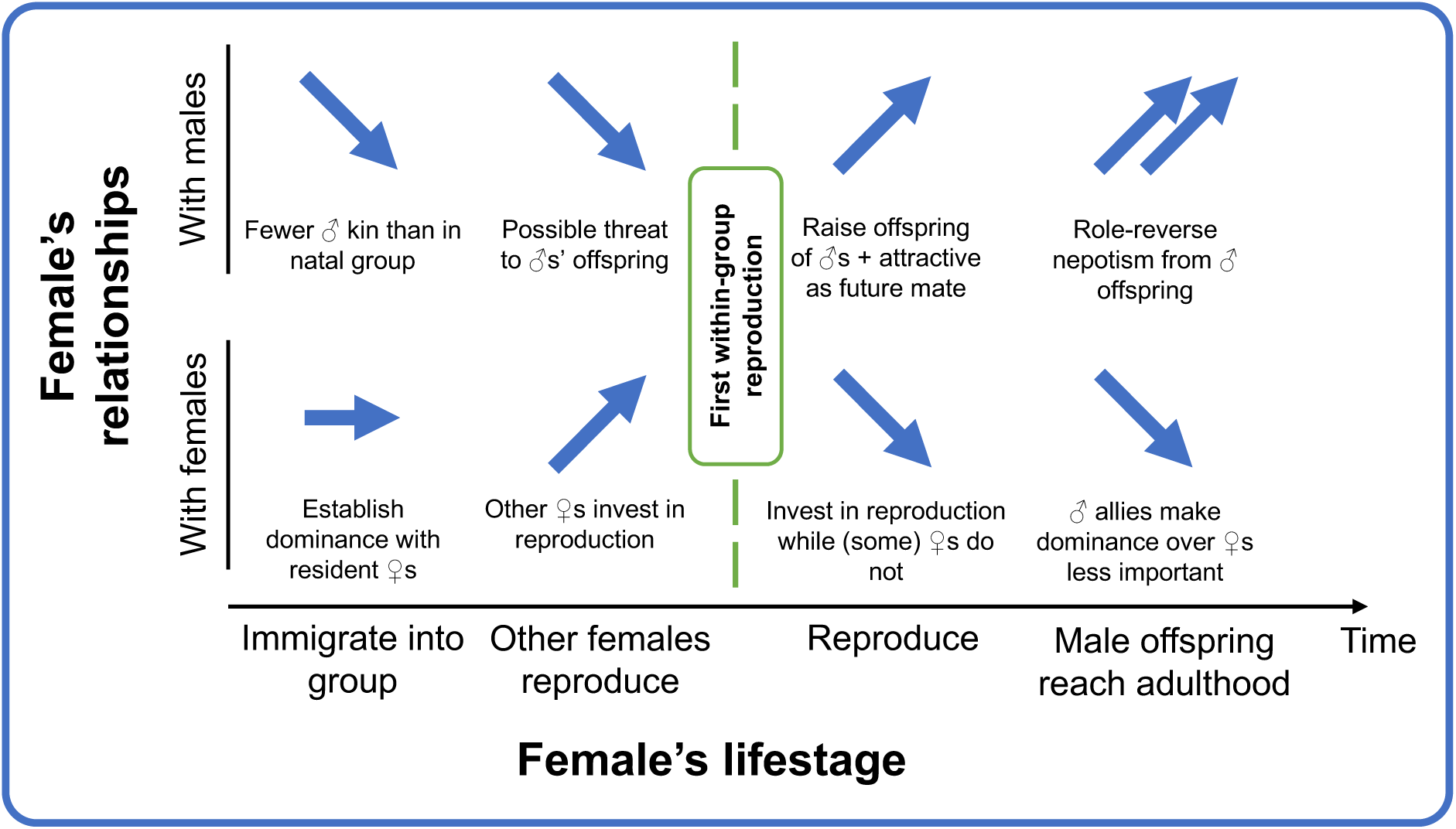
Schematic of females’ possible relationship transitions with both male and female adult group members over time—from immigrating into the social group through different reproductive events and, eventually, acquiring male within-group kin and associated benefits, as a consequence of reproduction. Arrows represent whether females’ relationships with males (affiliative) and females (agonistic) are proposed to improve, worsen, or remain unchanged during a particular lifestage.

### Considerations for dominance hierarchy inference

Our empirical results highlight how subtle variation in interaction rates among certain classes of group members can exist. As individuals and dyads in most animal groups vary in aspects such as kinship, pair bonding or reproductive status (as in our empirical data), similar variation in agonistic interaction rates can be expected to exist in many group-living species—shaping individuals’ social experience alongside outcomes. For example, tolerance is defined by a lack or low rate of aggression [69] and modulates how individuals experience their social environments alongside dominance [65,69,70]. However, current approaches to studying dominance typically consider interaction outcomes only (but see research on dominance interaction strategies, e.g. [71]). Considering rates of agonistic interactions, alongside their outcomes and contexts, may therefore provide a more complete understanding of individuals’ social behaviour.

Subtle variation in interaction rates can also prove problematic for dominance inference. Most existing dominance inference methods construct a group-level hierarchy directly from dyadic interactions, from which dyadic relationships are often retrospectively inferred via relative hierarchy positions [32,49]. Such methods rely on a balanced representation of individuals (and dyads) in interaction data, yet our empirical data highlight that this is not always the case. Our agent-based models show that hierarchy positions inferred via randomised Elo rating [32] and I&SI [49] approaches—which build group-level hierarchies directly from interactions—may be susceptible to such variation. These approaches consistently overestimated and underestimated particular individuals’ hierarchy positions in relation to variation in interaction rates that was associated with individuals’ traits (breeding status in our model). While greater data depth somewhat reduced the degree of overestimation and underestimation, the data depth of the initial simulation (>100 interactions per individual) was already considerable relative to what is often collected for empirical datasets. Furthermore, the degree of overestimation of ‘breeding females’ was greater when comparing females’ hierarchy positions to the entire group (Figure S8A,C). This demonstrates that females may also be inferred as dominant to males that they are intrinsically subordinate to as a consequence of variation in interaction rates, and could influence estimates of sex-stratification [12]. Meanwhile, the Percolation and Conductance [50] method—which first builds from the interaction to the dyadic level and only then the hierarchy—was robust to such variation in interaction rates among group members (see Figure 3C,D), and performed well even when comparing across the entire hierarchy (Figure S8E,F). It is important to remember that dominance hierarchies are an emergent property [72] and while informative, summarising individuals’ dyadic dominance relationship into group-level hierarchy positions, individuals’ positions remains an artificial construct [73]— akin to summarising an individual’s dyadic association or grooming relationships at the group level by their positions in constructed association [74] or grooming [75] networks.

It is difficult to determine whether overestimation and underestimation of hierarchy positions from the former two methods reflect something biologically meaningful. Such overestimated and underestimated hierarchy positions could, for example, better reflect resource access, as individuals with overestimated positions receive reduced aggression (at least in our model) and may thus be tolerated at resources ahead of dyadically dominant conspecifics. What is clear is that the Percolation and Conductance method, which also infers dyadic relationships directly [50], can robustly summarise individuals’ dyadic dominance relationships at the group (i.e. hierarchy) level irrespective of variation in interaction rates. This method thus represents a promising tool for inferring dominance hierarchy positions in wild and captive settings when interaction rates vary, and thereby also a first step towards solving the problem that the absence of interactions poses to dominance inference [33]. Such dominance hierarchies may then also provide a first step in determining whether individuals’ resource access agrees or disagrees with their hierarchy positions. If disagreement exists, this would directly imply that additional, biologically important processes—such as non-random tolerance or alliances— exist, contributing to individuals’ access to resources and, ultimately, shaping their social positions [1,76].

## Conclusions

We show that reproduction has contrasting consequences for females’ dominance interactions with female and male group members in vulturine guineafowl. Our study highlights a dynamic relationship between reproduction and social dominance in group-living species, and adds reproductive status as a further driver of dominance dynamics [77–79]. Given our limited dataset, we encourage future research to identify tangible long-term effects of breeding-related intersexual tolerance. For example, one could relate individuals’ food consumption rates, as well as patterns of intrasexual dominance interactions, at monopolisable food patches to both reproductive status and the presence of tolerant dominant-sex group members. Our empirical findings and agent-based models demonstrate the importance of interaction rates in the study of dominance: adding an often-overlooked facet (alongside interaction outcomes) to individuals’ experience of their competitive social landscapes, as well as demonstrating that interaction rates may influence the accurate estimation of individuals’ dominance hierarchy positions. The latter, based on empirically-inspired agent-based models, highlights Percolation and Conductance [50] as a robust tool for estimating both individuals’ hierarchy positions and group-level characteristics (e.g. degree of sex-stratification [12]). With our study, we hope to stimulate more research concerning the patterns and dynamics of dominance interactions (not just their outcomes) and relationships both within the subordinate sex and between the sexes, areas that have historically been understudied but are increasingly recognized as important [12,51,52,80].

## Author contributions

TD conceived of the initial idea for the study, which was further developed with input from BN, NJB and DRF. WC collected ∼40% of the breeding-season analysis dataset and ∼30% of the male-female analysis dataset; BN and TD collected ∼15% and ∼8%, respectively, of both the breeding-season analysis dataset and male-female analysis dataset. All trapping events that contributed measurements were overseen by BN. TD led all data preparation, analysis and visualization, as well as table preparation, with input from BN, WC, NJB and DRF. DRF conceived the initial agent-based model idea, which was developed, implemented, run and analysed by TD. TD wrote the initial manuscript, which was reviewed and improved upon with input from BN, WC, NJB and DRF. DRF was responsible for project funding acquisition and administration.

## Ethics

All field methods were ethically reviewed and approved by the Max Planck Society Ethikrat Committee (2016_13/1) and the Animal Ethics Committee of the Australian National University (A2023/20 Farine).

## Acknowledgements

We thank the Mpala Research Centre for logistical support, Kenya Wildlife Service for authorization to undertake the research (permit KWS-0016-01-21), the National Commission for Science, Technology and Innovation of Kenya (NACOSTI/P/16/3706/6465), the Wildlife Research and Training Institute (WRTI-0286-03-23), the National Environment Management Authority (NEMA/AGR/68/2017) for permits to Kenyan resources, and the National Museums of Kenya, and especially Dr Njoroge, for their ongoing support of our project. In addition, we thank Mary Waithira and Janet Wangare Kariuki for considerable assistance in the field, as well as Monicah Wambui, Edel Awour, John Wanjala, Elizabeth Koli and Mina Ogino for further assistance. We also thank Alastair Wilson for discussion of statistical approaches. Finally, we thank Josh Arbon for supplying R code that facilitated both fitting of ASReml models to empirical data as well as hierarchy inference in our agent-based models.

## Data availability

Data and code for all empirical aspects of this work will be published upon acceptance. Data and code for simulations are publicly available at: https://github.com/TobitDehnen/IntRatesDomInference.

## References

1. Sosa S. 2016 The influence of gender, age, matriline and hierarchical rank on individual social position, role and interactional patterns in *Macaca sylvanus* at ‘La Forêt des Singes’: a multilevel social network approach. Front. Psychol. 7, 529. (doi:10.3389/fpsyg.2016.00529)

2. Ward A, Webster M. 2016 Sociality: the behaviour of group-living animals. Switzerland: Springer International Publishing. (doi:10.1007/978-3-319-28585-6)

3. Shizuka D, McDonald DB. 2015 The network motif architecture of dominance hierarchies. J. R. Soc. Interface 12, 20150080. (doi:10.1098/rsif.2015.0080)

4. Ellis L. 1995 Dominance and reproductive success among nonhuman animals: a cross-species comparison. Ethol. Sociobiol. 16, 257–333. (doi:10.1016/0162-3095(95)00050-U)

5. Speakman JR. 2008 The physiological costs of reproduction in small mammals. Philos. Trans. R. Soc. B Biol. Sci. 363, 375–398. (doi:10.1098/rstb.2007.2145)

6. Harshman LG, Zera AJ. 2007 The cost of reproduction: the devil in the details. Trends Ecol. Evol. 22, 80–86. (doi:10.1016/j.tree.2006.10.008)

7. Williams GC. 1966 Natural selection, the costs of reproduction, and a refinement of Lack’s principle. Am. Nat. 100, 687–690. (doi:10.1086/282461)

8. Briffa M, Elwood RW. 2004 Use of energy reserves in fighting hermit crabs. Proc. R. Soc. Lond. B Biol. Sci. 271, 373–379. (doi:10.1098/rspb.2003.2633)

9. Neat FC, Taylor AC, Huntingford FA. 1998 Proximate costs of fighting in male cichlid fish: the role of injuries and energy metabolism. Anim. Behav. 55, 875–882. (doi:10.1006/anbe.1997.0668)

10. Hamilton WD. 1964 The genetical evolution of social behaviour. I. J. Theor. Biol. 7, 1–16. (doi:10.1016/0022-5193(64)90038-4)

11. Konečná M, Roubová V, Wallner B, Lhota S. 2023 The effect of maternal status on time budget in female barbary macaques (*Macaca sylvanus*). Int. J. Primatol. (doi:10.1007/s10764-023-00360-z)

12. Kappeler PM et al. 2022 Sex and dominance: How to assess and interpret intersexual dominance relationships in mammalian societies. Front. Ecol. Evol. 10, 918773. (doi:10.3389/fevo.2022.918773)

13. Marty PR, Hodges K, Agil M, Engelhardt A. 2016 Determinants of immigration strategies in male crested macaques (*Macaca nigra*). Sci. Rep. 6, 32028. (doi:10.1038/srep32028)

14. Vullioud C, Davidian E, Wachter B, Rousset F, Courtiol A, Höner OP. 2019 Social support drives female dominance in the spotted hyaena. *Nat*. Ecol. Evol. 3, 71–76. (doi:10.1038/s41559-018-0718-9)

15. Ilany A, Akçay E. 2016 Social inheritance can explain the structure of animal social networks. Nat. Commun. 7, 1–10. (doi:10.1038/ncomms12084)

16. Ilany A, Holekamp KE, Akçay E. 2021 Rank-dependent social inheritance determines social network structure in spotted hyenas. Science 373, 348–352. (doi:10.1126/science.abc1966)

17. Goldenberg SZ, Douglas-Hamilton I, Wittemyer G. 2016 Vertical transmission of social roles drives resilience to poaching in elephant networks. Curr. Biol. 26, 75–79. (doi:10.1016/j.cub.2015.11.005)

18. de Waal FBM. 1996 Macaque social culture: development and perpetuation of affiliative networks. J. Comp. Psychol. 110, 147–154. (doi:10.1037/0735-7036.110.2.147)

19. Berman CM, Rasmussen KLR, Suomi SJ. 1997 Group size, infant development and social networks in free-ranging rhesus monkeys. Anim. Behav. 53, 405–421. (doi:10.1006/anbe.1996.0321)

20. Greenwood PJ. 1980 Mating systems, philopatry and dispersal in birds and mammals. Anim. Behav. 28, 1140–1162. (doi:10.1016/S0003-3472(80)80103-5)

21. Romano A, Liker A, Bazzi G, Ambrosini R, Møller AP, Rubolini D. 2022 Annual egg productivity predicts female-biased mortality in avian species. Evolution 76, 2553–2565. (doi:10.1111/evo.14623)

22. Papageorgiou D, Farine DR. 2020 Shared decision-making allows subordinates to lead when dominants monopolize resources. Sci. Adv. 6, eaba5881. (doi:10.1126/sciadv.aba5881)

23. Nyaguthii B, Dehnen T, Klarevas-Irby JA, Papageorgiou D, Kosgey J, Farine DR. 2023 Cooperative breeding in a plural breeder: the vulturine guineafowl (*Acryllium vulturinum*). bioRxiv, 10.1101/2022.11.23.517633. (doi:10.1101/2022.11.23.517633)

24. Klarevas-Irby JA, Wikelski M, Farine DR. 2021 Efficient movement strategies mitigate the energetic cost of dispersal. Ecol. Lett. 24, 1432–1442. (doi:10.1111/ele.13763)

25. Breitenbach RP, Meyer RK. 1959 Effect of incubation and brooding on fat, visceral weights and body weight of the hen pheasant (*Phasianus colchicus*). Poult. Sci. 38, 1014–1026. (doi:10.3382/ps.0381014)

26. Papageorgiou D, Christensen C, Gall GEC, Klarevas-Irby JA, Nyaguthii B, Couzin ID, Farine DR. 2019 The multilevel society of a small-brained bird. Curr. Biol. 29, R1120–R1121. (doi:10.1016/j.cub.2019.09.072)

27. Del Hoyo J, Elliott A, Sargatal J, editors. 1994 Order Galliformes: Family Numididae (Guineafowl). In Handbook for the birds of the world. Vol 2. New World Vultures to Guineafowl, pp. 554–567. Barcelona: Lynx Edicions.

28. Papageorgiou D, Rozen-Rechels D, Nyaguthii B, Farine DR. 2021 Seasonality impacts collective movements in a wild group-living bird. Mov. Ecol. 9, 38. (doi:10.1186/s40462-021-00271-9)

29. Dehnen T, Papageorgiou D, Nyaguthii B, Cherono W, Penndorf J, Boogert NJ, Farine DR. 2022 Costs dictate strategic investment in dominance interactions. Philos. Trans. R. Soc. B Biol. Sci. 377, 20200447. (doi:10.1098/rstb.2020.0447)

30. Elbin SB, Crowe TM, Graves HB. 1986 Reproductive behavior of helmeted guinea fowl (*Numida meleagris*): mating system and parental care. Appl. Anim. Behav. Sci. 16, 179–197. (doi:10.1016/0168-1591(86)90110-3)

31. Altmann J. 1974 Observational study of behavior: sampling methods. Behaviour 49, 227–266. (doi:10.1163/156853974X00534)

32. Sánchez-Tójar A, Schroeder J, Farine DR. 2018 A practical guide for inferring reliable dominance hierarchies and estimating their uncertainty. J. Anim. Ecol. 87, 594–608. (doi:10.1111/1365-2656.12776)

33. Strauss ED, Shizuka D. 2022 The dynamics of dominance: open questions, challenges and solutions. Philos. Trans. R. Soc. B Biol. Sci. 377, 20200445. (doi:10.1098/rstb.2020.0445)

34. Strauss ED. 2023 Demographic turnover can be a leading driver of hierarchy dynamics, and social inheritance modifies its effects. Philos. Trans. R. Soc. B Biol. Sci. 378, 20220308. (doi:10.1098/rstb.2022.0308)

35. Wilson AJ, Grimmer A, Rosenthal GG. 2013 Causes and consequences of contest outcome: aggressiveness, dominance and growth in the sheepshead swordtail, *Xiphophorus birchmanni*. Behav. Ecol. Sociobiol. 67, 1151–1161. (doi:10.1007/s00265-013-1540-7)

36. Lane SM, Wilson AJ, Briffa M. 2020 Analysis of direct and indirect genetic effects in fighting sea anemones. Behav. Ecol. 31, 540–547. (doi:10.1093/beheco/arz217)

37. Hartig F. 2022 DHARMa: residual diagnostics for hierarchical (multi-level/mixed) regression models. R package version 0.4.6.

38. Fox J, Weisberg S. 2019 An R Companion to Applied Regression. Third edition. Thousand Oaks CA: Sage. See https://socialsciences.mcmaster.ca/jfox/Books/Companion/.

39. R Core Team. 2021 R: A language and environment for statistical computing.

40. Lüdecke D. 2023 sjPlot: data visualization for statistics in social science. R package version 2.8.14.

41. Wickham H. 2016 ggplot2: elegant graphics for data analysis. New York: Springer-Verlag. See https://ggplot2.tidyverse.org.

42. Csardi G, Nepusz T. 2006 The igraph software package for complex network research. InterJournal Complex Systems, 1695.

43. Wilke CO. 2020 cowplot: streamlined plot theme and plot annotations for ‘ggplot2’. R package version 1.1.1.

44. Wilson AJ, Morrissey MB, Adams MJ, Walling CA, Guinness FE, Pemberton JM, Clutton-Brock TH, Kruuk LEB. 2011 Indirect genetics effects and evolutionary constraint: an analysis of social dominance in red deer, *Cervus elaphus*. J. Evol. Biol. 24, 772–783. (doi:10.1111/j.1420-9101.2010.02212.x)

45. van Paridon JP, Bolker BM, Alday P. 2023 lmerMultiMember: Multiple membership random effects.

46. Bates D, Mächler M, Bolker B, Walker S. 2015 Fitting linear mixed-effects models using lme4. J. Stat. Softw. 67, 1–48. (doi:10.18637/jss.v067.i01)

47. Brooks ME, Kristensen K, van Benthem KJ, Magnusson A, Berg CW, Nielsen A, Skaug HJ, Mächler M, Bolker BM. 2017 glmmTMB balances speed and flexibility among packages for zero-inflated generalized linear mixed modeling. R J. 9, 378. (doi:10.32614/RJ-2017-066)

48. Ogino M, Strauss ED, Farine DR. 2023 Challenges of mismatching timescales in longitudinal studies of collective behaviour. Philos. Trans. R. Soc. B Biol. Sci. 378, 20220064. (doi:10.1098/rstb.2022.0064)

49. De Vries H. 1998 Finding a dominance order most consistent with a linear hierarchy: a new procedure and review. Anim. Behav. 55, 827–843. (doi:10.1006/anbe.1997.0708)

50. McCowan B, Vandeleest J, Balasubramaniam K, Hsieh F, Nathman A, Beisner B. 2022 Measuring dominance certainty and assessing its impact on individual and societal health in a nonhuman primate model: a network approach. Philos. Trans. R. Soc. B Biol. Sci. 377, 20200438. (doi:10.1098/rstb.2020.0438)

51. Clutton-Brock TH, Huchard E. 2013 Social competition and selection in males and females. Philos. Trans. R. Soc. B Biol. Sci. 368, 20130074. (doi:10.1098/rstb.2013.0074)

52. Clutton-Brock TH, Huchard E. 2013 Social competition and its consequences in female mammals. J. Zool. 289, 151–171. (doi:10.1111/jzo.12023)

53. Clutton-Brock TH. 1984 Reproductive effort and terminal investment in iteroparous animals. Am. Nat. 123, 212–229. (doi:10.1086/284198)

54. Dehnen T, Arbon JJ, Farine DR, Boogert NJ. 2022 How feedback and feed-forward mechanisms link determinants of social dominance. Biol. Rev. 97, 1210–1230. (doi:10.1111/brv.12838)

55. Holekamp KE, Ogutu JO, Dublin HT, Frank LG, Smale L. 1993 Fission of a spotted hyena clan: consequences of prolonged female absenteeism and causes of female emigration. Ethology 93, 285–299. (doi:10.1111/j.1439-0310.1993.tb01210.x)

56. van der Marel A, Francis X, O’Connell CL, Estien CO, Carminito C, Moore VD, Lormand N, Kluever BM, Hobson EA. 2023 Perturbations highlight importance of social history in parakeet rank dynamics. Behav. Ecol., arad015. (doi:10.1093/beheco/arad015)

57. Mohamad R, Monge J-P, Goubault M. 2010 Can subjective resource value affect aggressiveness and contest outcome in parasitoid wasps? Anim. Behav. 80, 629–636. (doi:10.1016/j.anbehav.2010.06.022)

58. Stokkebo S, Hardy ICW. 2000 The importance of being gravid: egg load and contest outcome in a parasitoid wasp. Anim. Behav. 59, 1111–1118. (doi:10.1006/anbe.2000.1407)

59. Chamorro-Florescano IA, Favila ME, Macías-Ordóñez R. 2017 Contests over reproductive resources in female roller beetles: outcome predictors and sharing as an option. PLOS ONE 12, e0182931. (doi:10.1371/journal.pone.0182931)

60. Ellis S et al. 2022 Patterns and consequences of age-linked change in local relatedness in animal societies. *Nat*. Ecol. Evol. 6, 1766–1776. (doi:10.1038/s41559-022-01872-2)

61. Koenig WD, Dickinson JL. 2016 Cooperative breeding in vertebrates: studies of ecology, evolution, and behavior. Cambridge: Cambridge University Press.

62. Lukas D, Huchard E. 2019 The evolution of infanticide by females in mammals. Philos. Trans. R. Soc. B Biol. Sci. 374, 20180075. (doi:10.1098/rstb.2018.0075)

63. Croft DP, Weiss MN, Nielsen MLK, Grimes C, Cant MA, Ellis S, Franks DW, Johnstone RA. 2021 Kinship dynamics: patterns and consequences of changes in local relatedness. Proc. R. Soc. B Biol. Sci. 288, 20211129. (doi:10.1098/rspb.2021.1129)

64. Newton-Fisher NE. 2014 Roving females and patient males: a new perspective on the mating strategies of chimpanzees: roving females and patient males. Biol. Rev. 89, 356–374. (doi:10.1111/brv.12058)

65. Van Horn RC, Wahaj SA, Holekamp KE. 2004 Role-reversed nepotism among cubs and sires in the spotted hyena (*Crocuta crocuta*). Ethology 110, 413–426. (doi:10.1111/j.1439-0310.2004.00984.x)

66. Anderson CM. 1986 Female age: male preference and reproductive success in primates. Int. J. Primatol. 7, 305–326. (doi:10.1007/BF02736394)

67. Nichols HJ, Amos W, Cant MA, Bell MBV, Hodge SJ. 2010 Top males gain high reproductive success by guarding more successful females in a cooperatively breeding mongoose. Anim. Behav. 80, 649–657. (doi:10.1016/j.anbehav.2010.06.025)

68. Ebensperger LA, Hurtado MJ. 2005 Seasonal changes in the time budget of degus, *Octodon degus*. Behaviour 142, 91–112. (doi:10.1163/1568539053627703)

69. Borgeaud C, Bshary R. 2015 Wild vervet monkeys trade tolerance and specific coalitionary support for grooming in experimentally induced conflicts. Curr. Biol. 25, 3011–3016. (doi:10.1016/j.cub.2015.10.016)

70. Chiarati E, Canestrari D, Vila M, Vera R, Baglione V. 2011 Nepotistic access to food resources in cooperatively breeding carrion crows. Behav. Ecol. Sociobiol. 65, 1791– 1800. (doi:10.1007/s00265-011-1187-1)

71. Hobson EA, Mønster D, DeDeo S. 2021 Aggression heuristics underlie animal dominance hierarchies and provide evidence of group-level social information. Proc. Natl. Acad. Sci. 118, e2022912118. (doi:10.1073/pnas.2022912118)

72. Aureli F, Schino G. 2019 Social complexity from within: how individuals experience the structure and organization of their groups. Behav. Ecol. Sociobiol. 73, 6. (doi:10.1007/s00265-018-2604-5)

73. Farine DR, Whitehead H. 2015 Constructing, conducting and interpreting animal social network analysis. J. Anim. Ecol. 84, 1144–1163. (doi:10.1111/1365-2656.12418)

74. Boogert NJ, Farine DR, Spencer KA. 2014 Developmental stress predicts social network position. Biol. Lett. 10, 1–5. (doi:10.1098/rsbl.2014.0561)

75. Borgeaud C, Sosa S, Sueur C, Bshary R. 2017 The influence of demographic variation on social network stability in wild vervet monkeys. Anim. Behav. 134, 155–165. (doi:10.1016/j.anbehav.2017.09.028)

76. Kawai M. 1958 On the rank system in a natural group of Japanese monkey (I). Primates 1, 111–130. (doi:10.1007/BF01813699)

77. Strauss ED, Holekamp KE. 2019 Social alliances improve rank and fitness in convention-based societies. Proc. Natl. Acad. Sci. 116, 8919–8924. (doi:10.1073/pnas.1810384116)

78. Combes SL, Altmann J. 2001 Status change during adulthood: life–history by–product or kin selection based on reproductive value? Proc. R. Soc. Lond. B Biol. Sci. 268, 1367– 1373. (doi:10.1098/rspb.2001.1631)

79. Portugal SJ, Usherwood JR, White CR, Sankey DWE, Wilson AM. 2020 Artificial mass loading disrupts stable social order in pigeon dominance hierarchies. Biol. Lett. 16, 20200468. (doi:10.1098/rsbl.2020.0468)

80. Davidian E, Surbeck M, Lukas D, Kappeler PM, Huchard E. 2022 The eco-evolutionary landscape of power relationships between males and females. Trends Ecol. Evol. 37, 706–718. (doi:10.1016/j.tree.2022.04.004)

