## Supplementary Materials for "Breeding alters females’ social positions by changing dominance dynamics"

### Title

#### Supplementary Materials

This PDF includes:

- Supplementary Text
- Supplementary Figures S1–S8
- Supplementary Tables S1–S3

#### Supplementary Text

**Post-breeding analysis**

In observational studies such as ours, breeders and non-breeders are unlikely to be random subsets of group members [1]. Therefore, uncontrolled forces could drive observed patterns. For example, individuals with increasing or declining dominance trajectories could choose to breed and generate any apparent effects. While nest predation acts to disrupt any mechanism segregating individuals into breeders and non-breeders in our study, this is unlikely to completely nullify such effects. We thus repeated our breeding-season analysis after the breeding period had finished in a follow-up analysis. Under the hypothesis that individuals with increasing or declining dominance trajectories choose to breed, we predicted that results of this post-breeding analysis would match those of the breeding-season analysis. For this post-breeding analysis, we created a dataset that emulates that of the breeding-season analysis, described above. Specifically, we created a seven-week interaction data subset immediately after breeding finished—approximately matching the duration of pre-breeding datasets in the breeding-season analysis. We then left a four-week gap, matching the breeding periods, and then created a second seven-week data subset—emulating the post-breeding data in the breeding-season analysis (Figure S1). For each season, each individuals' 'breeder' and 'non-breeder' status was kept from the prior breeding season. Thus, the dataset for this analysis was very similar to that of the breeding-season analysis: 1098 female-female interactions from 98 dyads, with the number of breeders in both seasons matching those of the breeding-season analysis. The model fitted for the post-breeding analysis was identical to that of the breeding-season analysis.

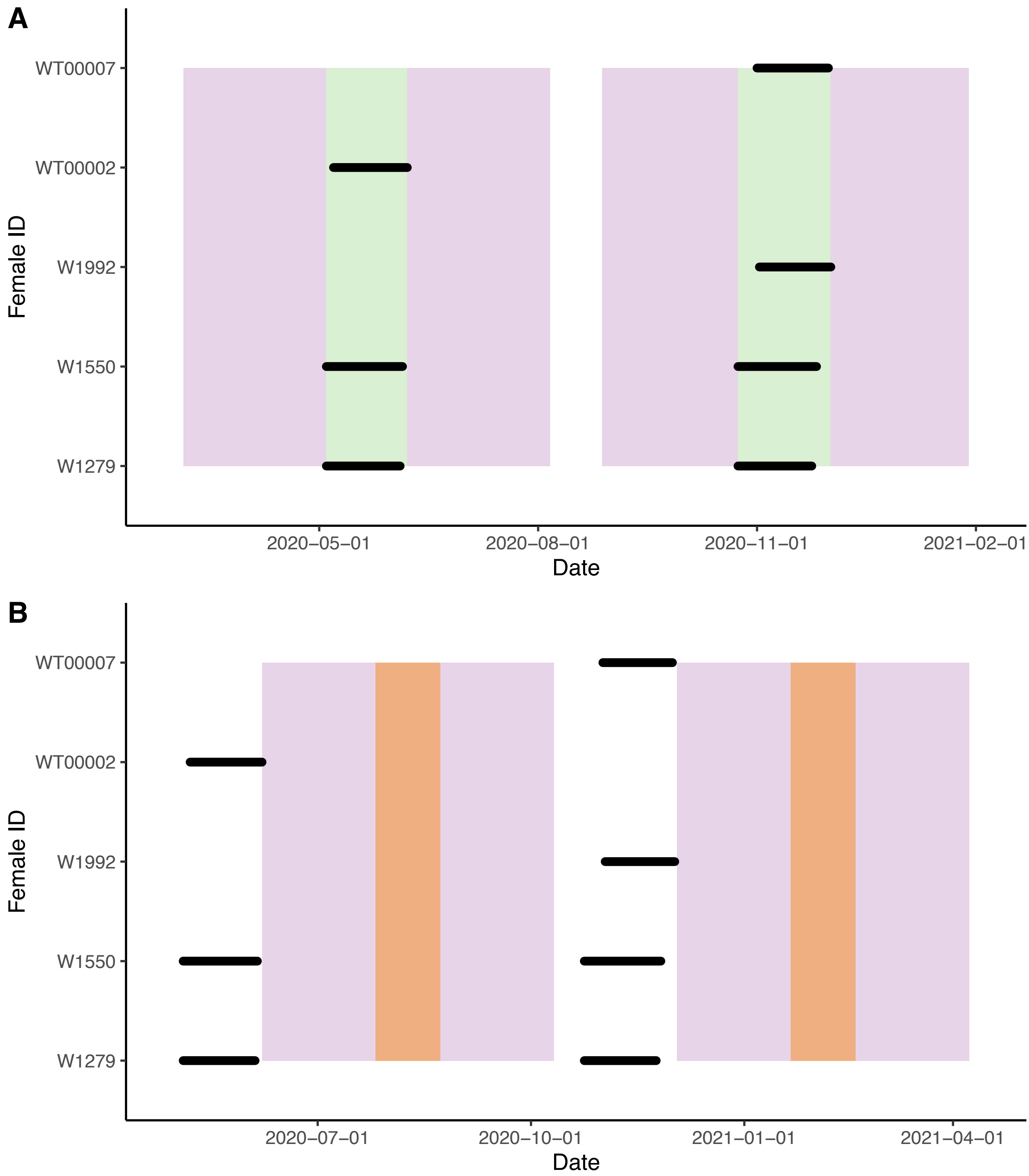


**Figure S1. Timeline of interaction data that contributed to breeding-related analyses.** Interaction data (purple blocks) are centralised around the halfway point of the breeding period (green blocks) in each season used in the breeding-season analysis (A). The post-breeding analysis is based on interaction data from two seven-week periods (purple blocks) that are separated by a four-week gap (orange blocks) for each season (B), thus emulating the breeding-season analysis in data structure. Horizontal black lines show the breeding period of each individual that hatched chicks in either season.

In this post-breeding analysis, we found no evidence that focals' winning probabilities differed between periods ($\chi^2_1$ = 1.516, P = 0.218) or seasons ($\chi^2_1$ = 1.314, P = 0.252). Rather, dyadic breeding contrast—i.e. the combination of the two interacting individuals' breeding status—drove much of the variation in predicted winning probability ($\chi^2_2$ = 33.608, p < 0.001): non-breeders were highly likely to win against breeders, breeders were highly likely to lose to non-breeders, and individuals of the same breeding category were similarly likely to win (Figure S2). We found no evidence of an interaction between period and dyadic breeding contrast ($\chi^2_2$ = 0.966, P = 0.617; Table S1), suggesting that results of the breeding-season analysis (Figure 2) are not driven by breeders being in a consistent, longitudinal decline. However, we note that the nature of binomial models constrains winning probability to lie between zero and one, and values are already relatively extreme for between-breeding category interactions after breeding. There is thus limited parameter space for the interaction term to be significant and in the same direction as in the breeding-season analysis. This result also highlights that the effect of breeding on winning probability is relatively persistent (the mid-points of the two periods in this analysis are approximately three months apart).

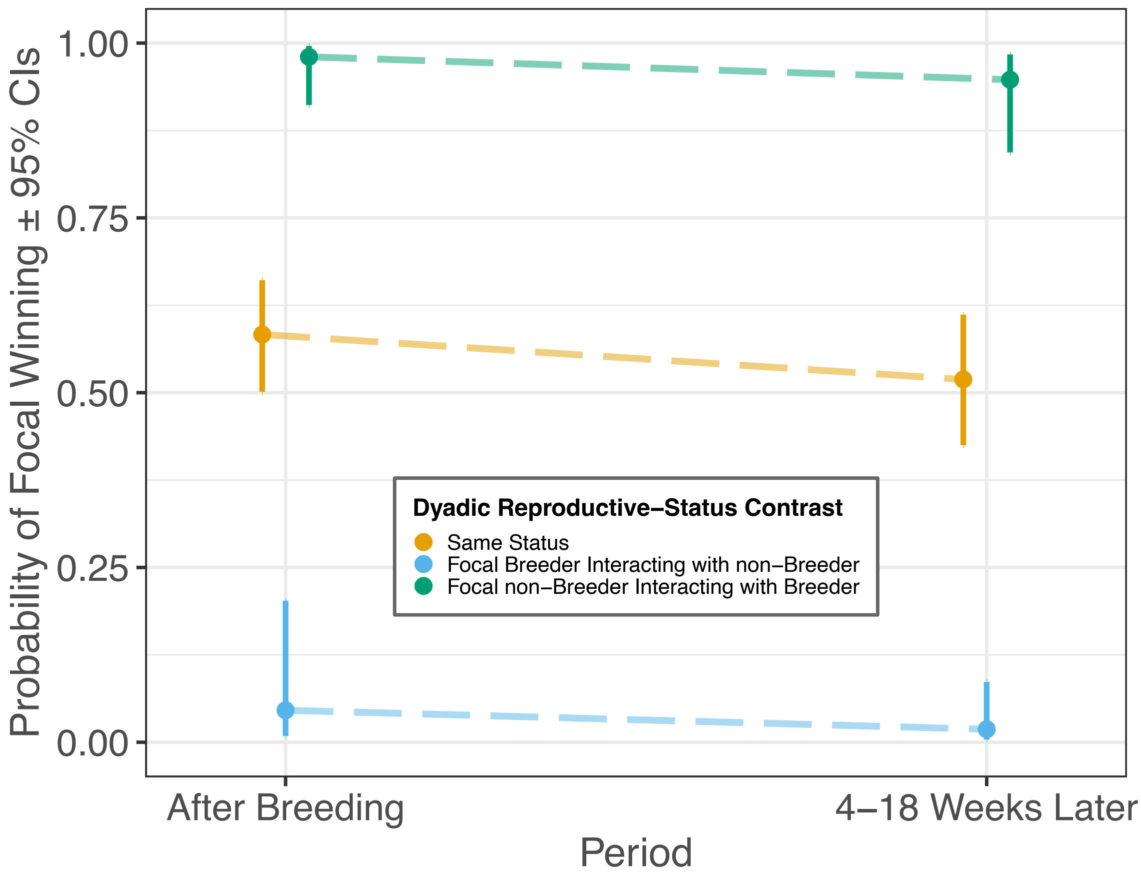


**Figure S2. Breeding females' probability of winning dominance interactions with non-breeding females remains low between two seven-week post-breeding periods separated by four weeks** (matching the breeding-season analysis in both dataset and model structure). Dyads where the focal and interacting individual are of the same breeding status (yellow points) comprise either a focal breeder interacting with another breeder or a focal non-breeder interacting with another non-breeder. Vertical bars represent 95% confidence intervals.

**Table S1. Predictors of winning probability among previously breeding and non-breeding female vulturine guineafowl.** The analysis matches the breeding-season analysis in the main text, replicated after breeding.

| Effect | $\chi^2$ | *d.f.* | *p value* |
|---|---|---|---|
| (Intercept) | 3.95 | 1 | **0.047** |
| period | 1.52 | 1 | 0.218 |
| dyadic_breeding_contrast | 33.61 | 2 | **<0.001** |
| season | 1.31 | 1 | 0.252 |
| period:dyadic_breeding_contrast | 0.97 | 2 | 0.617 |

Significant fixed effects are highlighted in bold.

**Comparison of lmerMultiMember and ASReml**

To ensure the robustness of our analytical methods, we fitted the breeding-season analysis model using the ASReml R package. This package has been used to control for individuals' IDs in many previous studies of contest outcomes where each individual contributes to multiple contest outcomes [2–5]. ASReml has the limitation of only fitting Gaussian error structures. However, while our data were not Gaussian, linear mixed-effects models are remarkably robust to violations of assumptions [6], and residual plots suggested that a linear mixed-effects model was not unreasonable. We thus created a replicate of the breeding-season analysis using the ASReml software and a Gaussian distribution. In this ASReml model we constrained the within-individual correlation between the focal_ID and interactor_ID random effects to be -1 [3], practically producing a single random term that encapsulates the additive effect of the focal and interactor IDs (for a comprehensive explanation see [4]). This replicates the random effects structure in the breeding-season analysis in the main text. To facilitate comparison, we also created an identical replicate of the breeding-season analysis using the lmerMultiMember model but with a Gaussian error distribution, such that only one factor (error distribution or software package) was changed between the models being compared. We found that the summaries of the ASReml and Gaussian lmerMultiMember models were extremely similar (see Tables S2 & S3). Furthermore, model predicted values for each level of the interaction term were nearly identical between the two models (Figures S3 & S4). We thus concluded that models fitted with these two software packages yield qualitatively similar results. Lastly, the predicted values from the Gaussian lmerMultiMember model (Figure S3) show the same trend as that of the binomial lmerMultiMember model (Figure 2), with fixed effect results from the two broadly matching.

**Table S2. Model summary of breeding-season analysis replicated using a Gaussian error distribution.** Predictors of focal winning probability among female vulturine guineafowl of

| Level | *Estimate* | *Std. Error* | *t-value* |
|---|---|---|---|
| (Intercept) | 0.50 | 0.02 | **20.84** |
| periodAfter Breeding | 0.00 | 0.03 | 0.03 |
| dyadic_breeding_contrastB->NB | -0.07 | 0.05 | -1.38 |
| dyadic_breeding_contrastNB->B | 0.04 | 0.05 | 0.81 |
| seasontwo | -0.01 | 0.02 | -0.33 |
| periodAfter Breeding:dyadic_breeding_contrastB->NB | -0.26 | 0.08 | **-3.41** |
| periodAfter Breeding:dyadic_breeding_contrastNB->B | 0.27 | 0.07 | **3.92** |

different breeding status (dyadic breeding contrast) before and after breeding (period) in two seasons from a linear mixed-effects model fitted using lmerMultiMember.
Model structure and reference levels are identical to the breeding-season analysis in the main text. Levels with an absolute t-value greater than 2 are highlighted in bold.

**Table S3. Summary of model identical in structure to that in Table S2, but fitted using ASReml.**

| Level | *Estimate* | *Std. Error* | *z-ratio* |
|---|---|---|---|
| (Intercept) | 0.50 | 0.02 | **20.81** |
| period_After Breeding | 0.00 | 0.03 | 0.03 |
| dyadic_breeding_contrast_B->NB | -0.07 | 0.05 | -1.37 |
| dyadic_breeding_contrast_NB->B | 0.04 | 0.05 | 0.81 |
| season_two | -0.01 | 0.02 | -0.33 |
| period_After Breeding:dyadic_breeding_contrast_B->NB | -0.26 | 0.08 | **-3.41** |
| period_After Breeding:dyadic_breeding_contrast_NB->B | 0.27 | 0.07 | **3.92** |

Model structure and reference levels are identical to the breeding-season analysis in the main text. Levels with an absolute z-ratio greater than 2 are highlighted in bold.

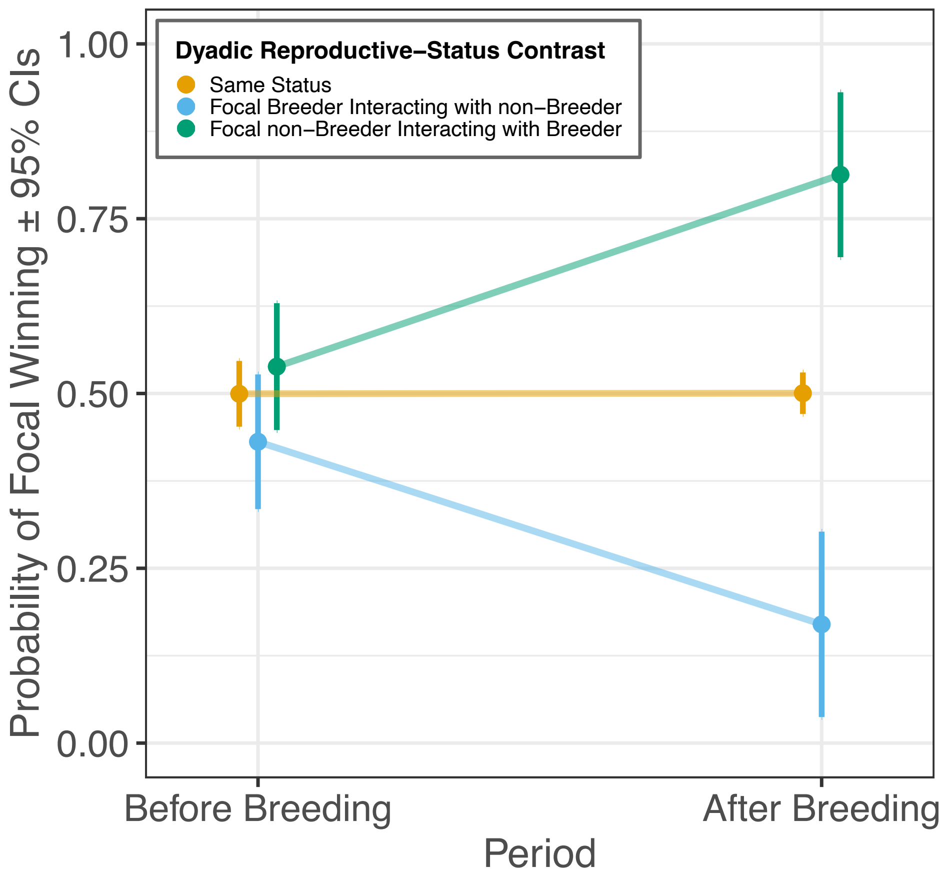


**Figure S3. Replicate of the breeding-season analysis but using a Gaussian error distribution.** As in the breeding-season analysis, the model was created either using the lmerMultiMember R package, differing from the breeding-season analysis only in the error structure.

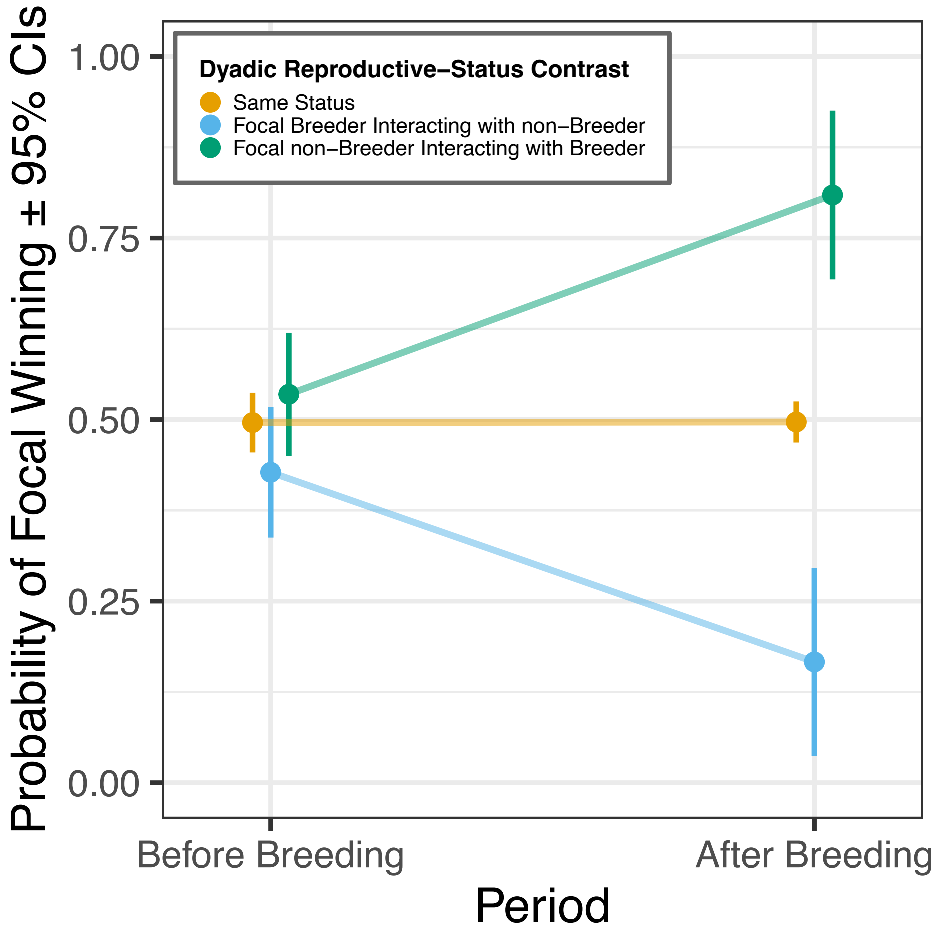


**Figure S4. Replicate of the breeding-season analysis but using a Gaussian error distribution and created in the R package ASReml.** The model thus differs to that used in Figure S3 only in the software used—but not in model structure.

**Long-term consequences of breeding: role-reversed nepotism**

Role-reversed nepotism is the phenomenon whereby adult individuals are more tolerant, or less aggressive, towards their parent(s) than equivalent non-parent group members [7]. Given that vulturine guineafowl females remain in social groups for years after reproducing, and males are both philopatric [8] and dominant over females [9], there is ample opportunity for role-reversed nepotism from adult males towards their mothers. To test for such role-reversed nepotism we used data on male-to-female aggressive interactions from 13$^{th}$ September 2019 until 9$^{th}$ March 2023. We removed data that were within one week of adult-offspring caring breeding interactions in order to focus the analysis on non-breeding periods, thereby excluding males' behaviour towards current breeders and minimising overlap with the short-term, male-female aggression analysis. For each data collection session longer than 15 minutes (to allow time for individuals to interact), we identified all male-female dyads for which the male's social mother—defined as the female that provided the vast majority of female offspring care to a given individual—was known and the male was older than 18 months. For each such male-female dyad in each data collection session, we identified whether the female was the social mother or not and whether the male had directed any aggression (1) or not (0) towards that female. We also calculated the proportion of the total interactions in that data collection session that took place at experimental food patches given that competition, and rates of associated agonistic interactions, should be heightened in such instances. Only data collection sessions where mother-son and non-mother-male dyads were present were included, as other sessions are not informative when including the data collection session as a random effect. This analysis was preliminary, due to drought-related lack of regular reproduction as well as high rates of juvenile predation resulting in only a single female producing male offspring that survived to adulthood. The dataset thus comprised a total of eight males and 25 females across 276 data collection sessions, with all mother-son dyads involving the same female. This analysis relied on dyadic data for each data collection session and did not require additive random effects structures due to all adult males being socially dominant to all females. We fitted a generalised linear mixed model with a binomial error distribution using the R package lme4 [10]. We fitted the binary variable male_aggressed_female as the response variable. Fixed effect included social_mother_or_not (whether the female was the social mother) the prop_ints_at_ExpPatch (the proportion of interactions that took place at an experimental patch, which was rescaled to aid model fitting) and their interaction term. Random effects included Group_ID (the data collection session), male_ID.original (male's ID) and female_ID.original (female's ID).

We found that males were less likely to aggress their mother than other female group members ($\chi^2_1$ = 5.632, P = 0.018; Figure S5A), consistent with the hypothesis that vulturine guineafowl males exhibit role-reversed nepotism towards their social mother. Males were also more likely to aggress females at experimental food patches ($\chi^2_1$ = 12.402, P < 0.001; Figure S5B). While model predicted values are relatively low (Figure S5), we note that these data were collected via all-occurrence sampling (and thus inevitably will omit some interactions that did occur) and during potentially short observation periods (>15 minutes) that are only a snapshot of the time individuals spent together. Thus, while aggression probabilities appear low, the observed pattern likely scales up to higher probabilities of dyadic aggressions that individual female vulturine guineafowl experience on a daily level.

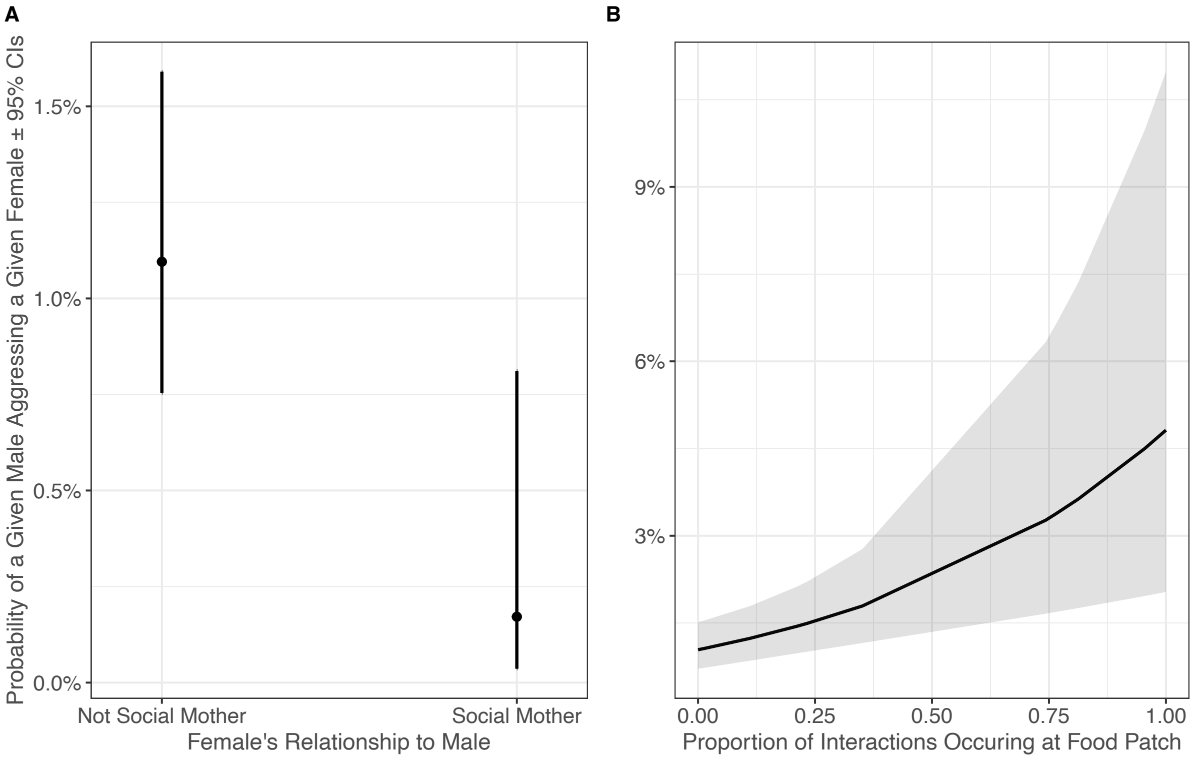


**Figure S5. Probability of a male vulturine guineafowl aggressing a female during all-occurrence sampling sessions (duration >15 minutes) as a function of (A) whether the female is the male's social mother or not and (B) the proportion of a group's interactions taking place at an experimental food patch.**

**Trapping analysis**

Results from our breeding-season analysis could be driven by various factors. For example, if breeders suffer reduced dominance this could be via reduced condition or their prolonged absence from the group. We thus asked whether there are physical predictors of intrasexual dominance interaction outcomes more generally, using interaction data outside the breeding season that surround trapping events where individuals' sizes and weights are known. Under the hypothesis that intrinsic attributes are important determinants of dominance among female vulturine guineafowl, we predicted that individuals that are structurally larger and in better condition would be more likely to win intrasexual dominance interactions.

For this analysis, we used body weight and tarsus length data from two whole group trapping events (on $9^{th}$ April 2021 and $9^{th}$ September 2022) of the focal social group used in the other analyses. We used these "snapshot" data in combination with interaction data from the trapping date ±2 weeks, comprising 498 male-male and female-female dominance interactions. We assumed individuals' tarsus and weight measurements to be unchanged across the four-week period. While this is unlikely to be strictly true, at least for weight, separate weight measurements from baited scales suggest that the weights of two adults and one female changed less than 4% between two weighing events separated by 27 days. Thus, given that the interaction data are at most two weeks from the trapping date, unaccounted variation in individuals' weights is likely minimal.

This analysis was conducted at the interaction level and thus the model used was similar in structure to the breeding-season and post-breeding analyses. The model contained only intrasexual interactions. In addition to the focal_ID and interactor_ID variables, which were generated by randomly allocating focal and interactor roles, we generated the following variables: focal_won, whether the focal won (1) or lost (0) the interaction; dyadic_sex_contrast, either M->M or F->F; dyadic_tarsus_contrast, calculated as focal tarsus length – interactor tarsus length; dyadic_weight_d_tarsus_contrast, calculated as (focal body weight / focal tarsus length) – (interactor body weight / interactor tarsus length); and trap_event, which trapping event the interaction took place in (one or two). We fitted a generalised linear mixed effects model with a binomial error structure using the lmerMultiMember R package [11]. The model had the following structure: our response variable was focal_won. We fitted dyadic_sex_contrast, dyadic_tarsus_contrast, and dyadic_weight_d_tarsus_contrast as fixed effects. As the effects of structural size and condition may be sex-specific, we also included an interaction term between dyadic_sex_contrast and dyadic_tarsus_contrast as well as dyadic_sex_contrast and dyadic_weight_d_tarsus_contrast. We wanted to control for trapping event (trap_event), which was fitted as a fixed effect due to having only two levels. As in the breeding-season and post-breeding analyses, we fitted focal_ID and interactor_ID as additive random effects, with Dyad_ID as a further random effect. We initially wanted to include an estimate of individuals' minimum age, as well as an interaction term between sex and the minimum age estimate, in our model. However, estimates of minimum age were correlated with condition, and thus omitted from the final model. Inspection of residuals suggests that model assumptions were met.

We found no difference in focal winning probability between interactions among males and those among females ($\chi^2_1$ = 0.742, P = 0.389). Sex was included as a fixed effect to allow for sex to be included in interaction terms but, as focal and interactor roles were allocated at random and all dominance interactions modelled here were *intrasexual*, the effect of sex itself was constrained to be insignificant by the structure of the data and model. Accordingly, this result does not suggest that sex differences in dominance ability are absent; in fact, before filtering the dataset for intrasexual dominance interactions, each of the 163 intersexual, male-female dominance interactions was won by the males—in keeping with previous data indicating that all adult males are dominant to females [9]. We found that structurally larger individuals—those with a longer tarsus—were more likely to win dominance interactions ($\chi^2_1$ = 4.788, P = 0.029; Figure S6A), and no evidence for this effect of structural size on winning probability to differ between male-male and female-female interactions ($\chi^2_1$ = 2.019, P = 0.155). There was a weak trend of individuals in better condition being more likely to win dominance interactions ($\chi^2_1$ = 3.015, P = 0.083; Figure S6B), and we again found no evidence for the effect of condition on winning probability to differ between male-male and female-female interactions ($\chi^2_1$ = 0.0358, P = 0.850).

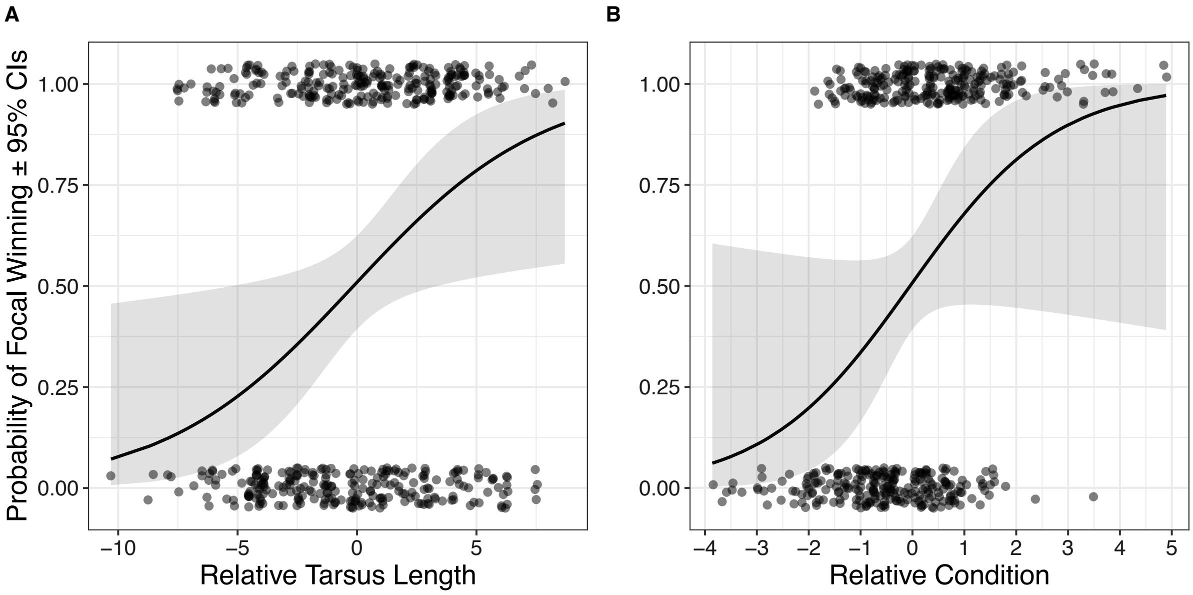


**Figure S6. Winning probability of focal individuals depending on (A) relative tarsus length and (B) relative condition.** Relative tarsus length was calculated as focal tarsus length minus interactor tarsus length, and relative condition was calculated as focal's body weight/tarsus length minus interactor's body weight/tarsus length. Black lines represent model predicted values while points represent the raw data. Note that raw data are binary and modelled and jittered for illustrative purposes only.

**The consequences of variation in interaction rates for dominance hierarchies**

Our empirical results suggest that, after breeding, breeding females receive less male aggression than non-breeding females (Figure 2). We simulated the consequences of such variation in interaction rates on females' inferred hierarchy positions using agent-based models as follows.

1. We generated a social group comprising 16 males and 10 females.
2. We created two normal distributions of dominance values, one with mean = 0 and standard deviation = 1 and a second with mean = -4 and standard deviation = 1; females were assigned the lowest 10 dominance values and males the highest 16 dominance values, such that all males were dominant to all females. We then standardised dominance values, such that each individual had a value between zero and 25. These dominance values were used to determine interaction outcomes, and thus the order of dominance values reflected individuals' 'real hierarchy positions'.
3. We randomly assigned a certain proportion of females as 'breeding' (representing breeding females in our empirical data) and all other females as 'non- breeding'—this was set to 50% in the main results.

4. We simulated the outcomes of interactions among male-male, male-female, and female-female dyads as follows. For an interaction among individuals A and B, the probability of individual A winning ($P_A$) was calculated as:

$$P_A = \frac{1}{1 + e^{-S(D_A - D_B)}}$$

   Where $D_A$ and $D_B$ represent the dominance values of individuals A and B, respectively, and $S$ is a steepness parameter—modulating the hierarchy steepness and set to a value of one for the main results. With a difference in dominance values of one and steepness parameter, $S$, of one, a dominant individual has a 73% probability of winning a given interaction. Whether individual A won was then calculated using the *sample* R function using the probability $P_A$.
5. Using this approach, we simulated intrasexual and intersexual interactions, with males generally engaging in interactions more so than females. We modelled two scenarios:
   i. An unbiased scenario, where males tolerate breeding and non-breeding females equally. Under this scenario we simulated 12 interactions for each male-male dyad, eight interactions for each male-female dyad and four interactions for each female-female dyad (total no. interactions = 2900).
   ii. A biased scenario, where males tolerate breeding females but not non-breeding females, which also influences female-female interactions. Under this scenario males redirect intersexual aggression away from breeding, i.e. tolerated, females and towards non-breeding females. Accordingly, males did not engage in interactions with breeding females and instead spread these interactions evenly among non-breeding females. The increase in the number of intersexual interactions each non-breeding female was involved in was thus directly, and inversely, proportional to the proportion of females that were designated as breeders: when 50% of females were breeders, each non-breeding female was involved in twice as many male-female interactions. In addition to males redirecting their intersexual interactions, males' tolerance of breeding females was modelled to influence within-female category interaction rates, such that dyads comprising breeding and non-breeding females engaged in half the number of interactions as in the unbiased scenario. Thus, breeding females experienced no male aggression and, via male tolerance—such as may occur on food patches—that was not granted to non-breeding females, engaged in fewer intrasexual agonistic interactions in this scenario (total no. interactions = 2850).
6. For each scenario (i and ii in step five) we then: inferred a dominance hierarchy and resulting female-only rank orders from simulated interaction outcomes using one of three methods (see below), calculated the difference between the inferred hierarchy

position and real hierarchy position for each female, and then calculated the mean difference for both breeding and non-breeding females.

We repeated steps one to six 1000 times in the main analysis (500 times for parameter exploration), and plotted density distributions, as well as 15, 50 and 85% quantiles, for preferred and non-preferred females' mean values from each run for each scenario. We repeated this process using three hierarchy inference methods. Specifically, we used randomised Elo ratings [12] and I&SI [13], two methods which build hierarchies directly from interactions, as well as Percolation and Conductance [14], which first builds dyadic dominance relationships from interaction data, and then uses the dyadic relationships to build the group-level dominance hierarchy. Lastly, we repeated this using entire group hierarchy rank orders; thus, if females were inferred as dominant to males this was also captured in the difference between real hierarchy positions and inferred hierarchy positions (see Figure S8).

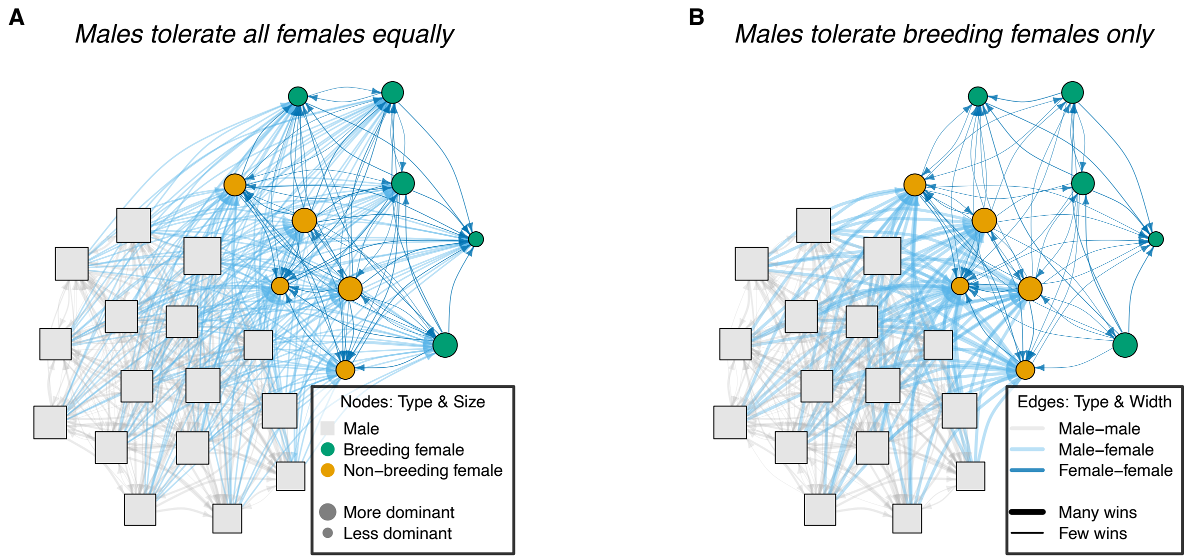


**Figure S7. Winner-loser networks of simulated agonistic interactions among males and females in two interaction scenarios** (used in the agent-based models of interaction rates and hierarchy inference). Males, which here interact more than females, either: (A) tolerate all females equally irrespective of their reproductive status (breeding or non-breeding); or (B) tolerate breeding females only, whereby males redirect intersexual interactions from breeding to non-breeding females with a concomitant reduction in the number of interactions between breeding vs non-breeding females. Edges are directed and point from winners to losers.

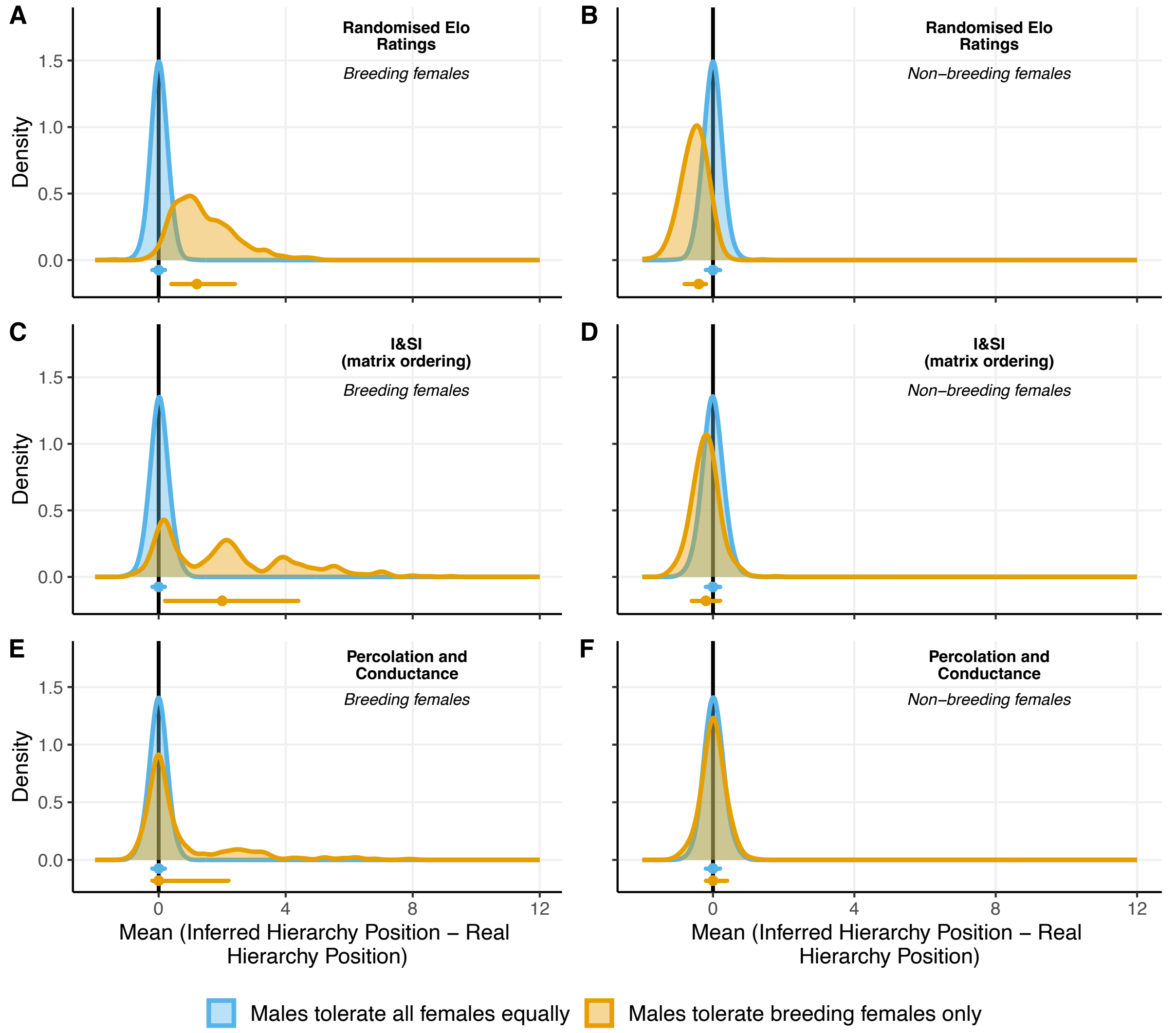


**Figure S8. Replicate of main agent-based model results (Figure 3) differing only in that female's inferred and real hierarchy positions were calculated at the group level (rather than only among females) and half as many simulations (500) were used here.** Note that axes here are scaled differently than in Figure 3.

#### References

1. Blount JD, Vitikainen EIK, Stott I, Cant MA. 2016 Oxidative shielding and the cost of reproduction. *Biol Rev* **91**, 483–497. (doi:10.1111/brv.12179)

2. Wilson AJ, Grimmer A, Rosenthal GG. 2013 Causes and consequences of contest outcome: aggressiveness, dominance and growth in the sheepshead swordtail, *Xiphophorus birchmanni*. *Behav Ecol Sociobiol* **67**, 1151–1161. (doi:10.1007/s00265-013-1540-7)

3. Lane SM, Wilson AJ, Briffa M. 2020 Analysis of direct and indirect genetic effects in fighting sea anemones. *Behavioral Ecology* **31**, 540–547. (doi:10.1093/beheco/arz217)

4. Wilson AJ, Morrissey MB, Adams MJ, Walling CA, Guinness FE, Pemberton JM, Clutton-Brock TH, Kruuk LEB. 2011 Indirect genetics effects and evolutionary constraint: an

analysis of social dominance in red deer, *Cervus elaphus*. *Journal of Evolutionary Biology* **24**, 772–783. (doi:10.1111/j.1420-9101.2010.02212.x)

5. Santostefano F, Wilson AJ, Araya-Ajoy YG, Dingemanse NJ. 2016 Interacting with the enemy: indirect effects of personality on conspecific aggression in crickets. *BEHECO* **27**, 1235–1246. (doi:10.1093/beheco/arw037)

6. Schielzeth H *et al.* 2020 Robustness of linear mixed-effects models to violations of distributional assumptions. *Methods Ecol Evol* **11**, 1141–1152. (doi:10.1111/2041-210X.13434)

7. Van Horn RC, Wahaj SA, Holekamp KE. 2004 Role-reversed nepotism among cubs and sires in the spotted hyena (*Crocuta crocuta*). *Ethology* **110**, 413–426. (doi:10.1111/j.1439-0310.2004.00984.x)

8. Klarevas-Irby JA, Wikelski M, Farine DR. 2021 Efficient movement strategies mitigate the energetic cost of dispersal. *Ecology Letters* **24**, 1432–1442. (doi:10.1111/ele.13763)

9. Papageorgiou D, Farine DR. 2020 Shared decision-making allows subordinates to lead when dominants monopolize resources. *Science advances* **6**, eaba5881. (doi:10.1126/sciadv.aba5881)

10. Bates D, Mächler M, Bolker B, Walker S. 2015 Fitting linear mixed-effects models using lme4. *Journal of Statistical Software* **67**, 1–48. (doi:10.18637/jss.v067.i01)

11. van Paridon JP, Bolker BM, Alday P. 2023 lmerMultiMember: Multiple membership random effects.

12. Sánchez-Tójar A, Schroeder J, Farine DR. 2018 A practical guide for inferring reliable dominance hierarchies and estimating their uncertainty. *Journal of Animal Ecology* **87**, 594–608. (doi:10.1111/1365-2656.12776)

13. De Vries H. 1998 Finding a dominance order most consistent with a linear hierarchy: a new procedure and review. *Animal Behaviour* **55**, 827–843. (doi:10.1006/anbe.1997.0708)

14. McCowan B, Vandeleest J, Balasubramaniam K, Hsieh F, Nathman A, Beisner B. 2022 Measuring dominance certainty and assessing its impact on individual and societal health in a nonhuman primate model: a network approach. *Phil. Trans. R. Soc. B* **377**, 20200438. (doi:10.1098/rstb.2020.0438)
